# Splicing buffers suboptimal codon usage in human cells

**DOI:** 10.1101/527440

**Authors:** Christine Mordstein, Rosina Savisaar, Robert S Young, Jeanne Bazile, Lana Talmane, Juliet Luft, Michael Liss, Martin S Taylor, Laurence D Hurst, Grzegorz Kudla

## Abstract

Although multiple studies have addressed the effects of codon usage on gene expression, such studies were typically performed in unspliced model genes. In the human genome, most genes undergo splicing and patterns of codon usage are splicing-dependent: guanine and cytosine (GC) content is highest within single-exon genes and within first exons of multi-exon genes. Intrigued by this observation, we measured the effects of splicing on expression in a panel of synonymous variants of GFP and mKate2 reporter genes that varied in nucleotide composition. We found that splicing promotes the expression of adenine and thymine (AT)-rich variants by increasing their steady-state protein and mRNA levels, in part through promoting cytoplasmic localization of mRNA. Splicing had little or no effect on the expression of GC-rich variants. In the absence of splicing, high GC content at the 5’ end, but not at the 3’ end of the coding sequence positively correlated with expression. Among endogenous human protein-coding transcripts, GC content has a more positive effect on various expression measures of unspliced, relative to spliced mRNAs. We propose that splicing promotes the expression of AT-rich genes, leading to selective pressure for the retention of introns in the human genome.

## Introduction

Mammalian genomes are characterised by large regional variation in base composition (Bernardi, 1993). Regions with a high density of G and C nucleotides (GC-rich regions) are in an open, transcriptionally active state, are gene-dense, and replicate early. In contrast, AT-rich regions are enriched with heterochromatin, contain large gene deserts and replicate late (Arhondakis et al., 2011; Lander et al., 2001; Vinogradov, 2003). The mechanisms that give rise to this compositional heterogeneity have been under debate for years and many researchers believe that the pattern originates from the process of GC-biased gene conversion (Duret and Galtier, 2009), though other neutral and selective mechanisms have been proposed as well (Eyre-Walker, 1991; Galtier et al., 2018; Plotkin and Kudla, 2011; Sharp and Li, 1987b).

The sequence composition of mammalian genes correlates with the GC-content of their genomic location. Thus, introns and exons of genes located in GC-rich parts of the genome are themselves GC-rich. This can potentially influence gene expression in multiple ways: nucleotide composition affects the physical properties of DNA, the thermodynamic stability of RNA folding, the propensity of RNA to interact with other RNAs and proteins, the codon adaptation of mRNA to tRNA pools, and the propensity for RNA modifications, such as m6A (Dominissini et al., 2012) and ac4C (Arango et al., 2018). Strikingly, studies of the effects of nucleotide composition on gene expression in human cells have led to opposing conclusions. On the one hand, heterologous expression experiments typically report large positive effects of GC content on protein production in a wide variety of transgenes, including fluorescent reporter genes, human cDNAs, and viral genes (Bauer et al., 2010; Kosovac et al., 2011; Kotsopoulou et al., 2000; Kudla et al., 2006; Zolotukhin et al., 1996). As a result, increasing the GC content of transgenes has become a common strategy in coding sequence optimization for heterologous expression in human cells (Fath et al., 2011). On the other hand, genome-wide analyses of endogenous genes typically show little or no correlation of GC content with expression (Duan et al., 2013; Lercher et al., 2003; Rudolph et al., 2016; Semon et al., 2005).

We hypothesized that the conflicting results in heterologous and endogenous gene expression studies can be partially explained by RNA splicing. Most transgenes used in heterologous expression systems have no introns, whereas 97% of genes in the human genome contain one or more introns. Splicing is known to influence gene expression at multiple stages, including nuclear RNP assembly, RNA export, and translation. If splicing selectively increased the expression of AT-rich genes, it could account for the lack of correlation of GC content and gene expression in previous genome-wide studies. We therefore compared spliced and unspliced genes with respect to their (1) genomic codon usage, (2) expression levels of reporter genes in transient and stable transfection experiments and (3) global expression patterns in human transcriptome studies. We show that splicing increases the expression of AT-rich genes, but not GC-rich genes, in part through effects on cytoplasmic RNA enrichment.

## Results

### Codon usage of human protein-coding genes depends on RNA splicing

We first analysed the relationship between the nucleotide composition of human genes and splicing. GC4 content (GC content at 4-fold degenerate sites) correlates negatively with the number of exons in humans (Figure 1A; Spearman’s ρ = −0.27; p < 2.2×10^−16^; see also (Carels and Bernardi, 2000; Ressayre et al., 2015; Savisaar and Hurst, 2016)). In addition, GC4 content is highest in 5’-proximal exons (Figure 1B; Spearman’s ρ = −0.18; p < 2.2×10^−16^), and first exons have a higher GC4 content than second exons (p < 2.2×10^−16^, one-tailed Wilcoxon test). Although these patterns could result from proximity to CpG-rich transcription start sites (TSSs)(Zhang et al., 2004), we found that first exons have significantly higher GC4 content than second exons even when controlling for the distance from the TSS (Figure 1C). This suggests that splicing contributes to the observed enrichment of G and C nucleotides in the 5’-proximal exons in human.

**Figure 1.**
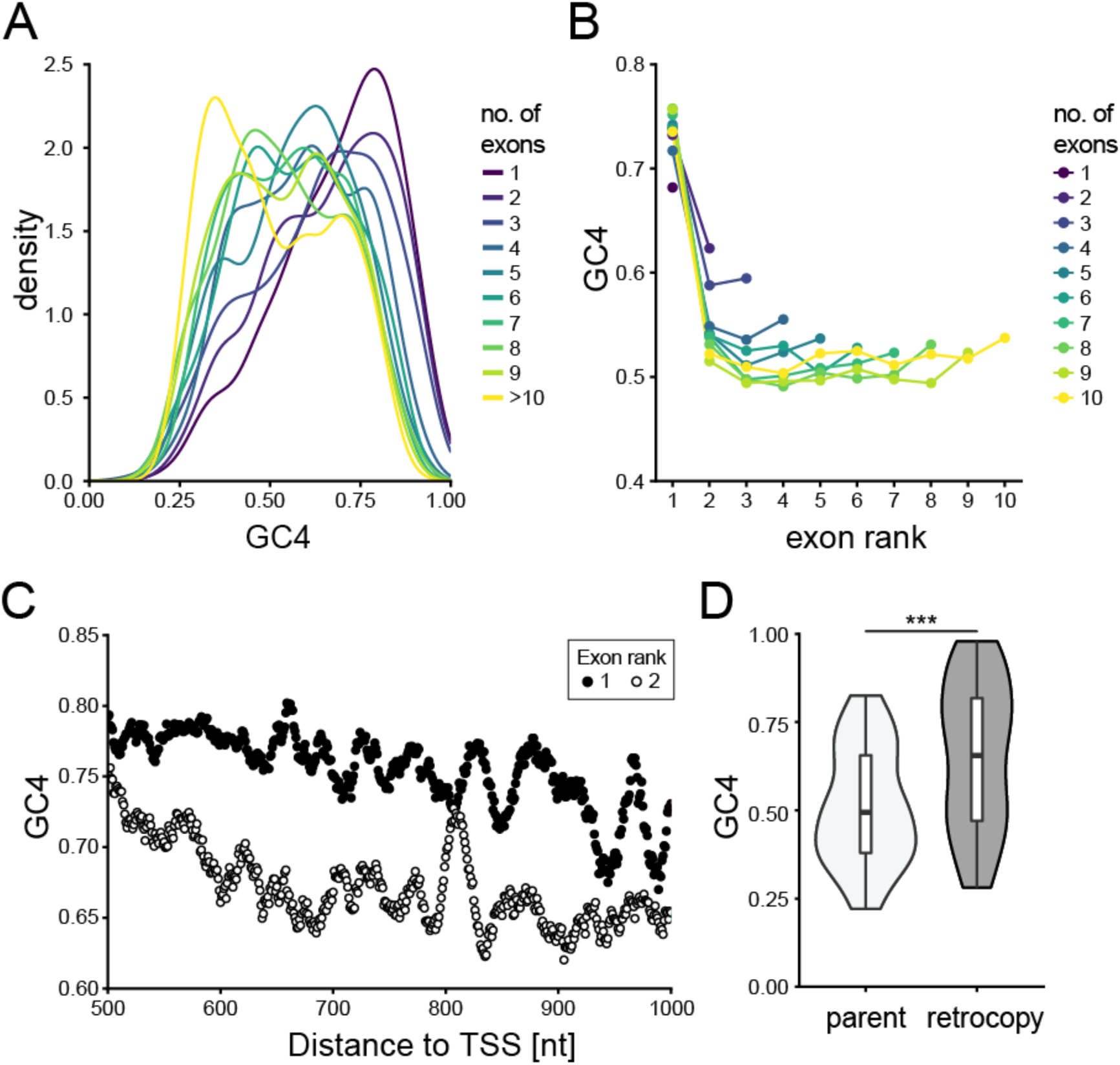
Splicing- and position-dependent patterns of nucleotide composition in human genes. (A) GC4 distribution of human protein-coding genes, grouped by number of exons per gene. (B) Mean GC4 content in protein-coding exons, grouped by exon position (rank) and by number of exons per gene. (C) Mean GC4 for individual codons within exons of rank 1 (black dots) or rank 2 (white dots) downstream of the transcription start site (TSS). (D) GC4 distribution of functional retrogenes (dark grey) and their corresponding parental genes (light grey) conserved between mouse and human (p=2.1×10^−4^, from one-tailed Wilcoxon signed rank test, n=49).

To understand the causal links between splicing and nucleotide composition, we studied the compositional patterns of retrogenes. Retrotransposition provides a natural evolutionary experiment of what happens when a previously spliced gene suddenly loses its introns. We first analysed a set of 49 parent-retrogene pairs for which both the parent and the retrocopy ORFs have been retained in human and mouse. Strikingly, we found that the retrocopies had a significantly higher GC4 content than their parents (median GC4_retrocopy_ - GC4_parent_ = 11.5%; p = 2.1×10^−4^ from one-tailed Wilcoxon test; Figure 1D). It thus appears that after retrotransposition, newly integrated intronless genes come under selective pressure for increased GC content. In a comparison of 31 parent-retrogene pairs retained between human and macaque, the median GC4 difference is not significant (0.09%; p = 0.13, Wilcoxon test), but this may be explained by duplication events in macaques being more recent (dS ~0.06) than in mouse (dS ~ 0.5) and therefore less evolutionary time has passed to allow changes in GC composition to have occurred. As a control, we analysed the GC4 content of retrocopies classed as pseudogenes (Figure S1A) and found it to be significantly lower compared to their parental genes (−2.963%; p < 2.2×10^−16^, Wilcoxon test). Furthermore, the genomic neighbourhood of functional retrocopies and pseudogenes had significantly lower GC content than the neighbourhood of their respective parental genes (Figure S1B). These observations suggest that increased GC content is not intrinsically connected with retrotransposition, but is required for maintaining long-term functionality of retrogenes. Taken together, these results support a splicing-dependent mechanism shaping conserved patterns of nucleotide composition across functional protein-coding genes.

### GC-content is a strong predictor of expression of unspliced reporter genes

The above analyses show a connection between splicing and genomic GC content of endogenous human genes. To test whether splicing differentially affects the expression of genes depending on their GC content, we designed 22 synonymous variants of GFP that span a broad range of GC3 content (GC content at the third positions of codons) (Mittal et al., 2018) (Figure S2). The collection encompasses most of the variation in GC3 content found among human genes. All variants were independently designed by randomly drawing each codon from an appropriate probability distribution, to ensure uniform GC content and statistical independence between sequences. We cloned these variants into two mammalian expression vectors: an intronless vector with a CMV promoter (pCM3) and a version of the same vector with a synthetic intron located in the 5’ UTR (pCM4). The GC content profiles of the 5’ UTRs were similar in both vectors (Figure S2E,F). The vectors also encoded a far-red fluorescent protein, mKate2, which we used to normalize GFP protein abundance (normalization reduced measurement noise, but similar results were obtained with and without normalization). Transient transfections of HeLa cells with three independent preparations of each plasmid showed reproducible expression with a large dynamic range: synonymous variants differed in GFP protein production 46-fold. Consistent with previous studies, GFP fluorescence was strongly correlated with GC3 content, both in spliced and unspliced genes (Figure 2A,B). Interestingly, introduction of an intron into the 5’ UTR increased the expression of most, but not all variants. Typically, GC-poor variants experienced a large increase of expression in the presence of an intron, whereas GC-rich variants were unaffected or experienced a moderate increase (Figure 2C).

**Figure 2.**
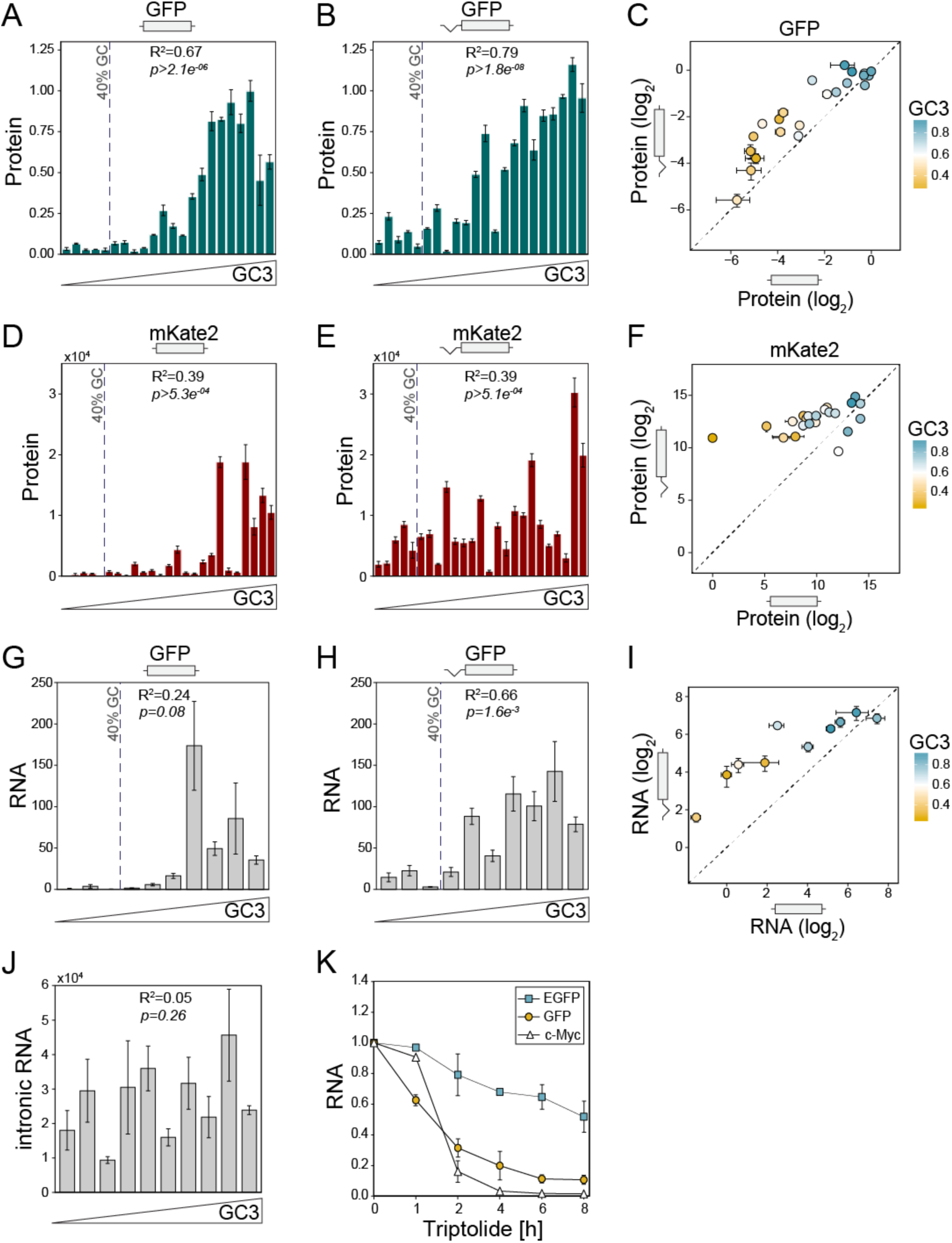
The effect of GC content on gene expression depends on splicing. (A-B) Protein levels of 22 GFP variants when transiently expressed as unspliced (A) or spliced (B) constructs were expressed in HeLa cells and quantified by spectrofluorometry. Each data point represents the mean of 9 replicates, +/-SEM. (C) Correlation of protein levels between unspliced and spliced variants of GFP (n=22, R^2^=0.69, p=9.0×10^−7^). The dashed line indicates x=y. (D-E) Protein levels of 23 mKate2 variants in the absence (A) or presence (B) of splicing. Each data point represents the mean of 9 replicates. (F) Correlation of protein levels between unspliced and spliced variants of mKate2 (n=23, R^2^=0.29, p=2.8×10^−4^). (G-H) mRNA levels of 10 GFP variants when transiently expressed as unspliced (G) or spliced (H) constructs in HeLa cells and quantified by qRT-PCR. Data points represent the mean of 3 replicates, +/-SEM. (I) Comparison of mRNA expression from spliced and unspliced GFP variants. (J) Intronic RNA levels of GFP variants measured by qRT-PCR. (K) RNA stability time course of GC-rich (97% GC3; t_1/2_=8.6h) and GC-poor (33% GC3, t_1/2_=2.4h) GFP variants in stably transfected HeLa Flp-In cells after blocking transcription with 500nM Triptolide. Results represent the averages of 2 independent experiments, +/-SD.

We obtained similar results in stably transfected HeLa and HEK293 cells (Figure S3) and when expressing an independently designed collection of 25 synonymous variants of mKate2 in HeLa cells (Figure 2D-F, S2D). A Fisher’s exact test revealed that the percentage of variants whose expression was increased by splicing significantly depended on GC3 content (p=0.02, N=47, GFP and mKate variants combined). These experiments show that many AT-rich genetic variants are expressed inefficiently in human cells, but low expression can be partially rescued by splicing. Notably, the average GC content of the human genome is 41% (Li, 2011). In our experiments, genes with GC content below 41% are expressed extremely inefficiently, unless they contain an intron (Figure 2). This may provide a strong selective pressure for the retention of introns in many human genes.

To establish which stages of expression are responsible for this phenomenon, we first measured mRNA abundance of GFP variants in transiently transfected HeLa cells by quantitative RT-PCR (qRT-PCR). High GC content may introduce unwanted bias in RT-PCR, so to allow fair comparison of all variants irrespective of their GC content, qRT-PCR primers were placed in the untranslated regions, whose sequence did not vary. Similar to protein levels, mRNA abundance varied widely between synonymous variants of GFP. GC-poor variants experienced a large increase of expression in the presence of an intron, whereas GC-rich variants were less affected (Figure 2G-I). The range of variation in mRNA abundance was much smaller in constructs with an intron than without intron (Figure 2I), indicating that splicing buffers the effects of GC content on expression.

We then asked if changes in mRNA abundance arose at transcriptional or post-transcriptional levels. As a proxy for transcriptional efficiency, we measured the abundance of intronic RNA for GFP variants expressed from the intron-containing plasmid. GC content did not correlate with intronic RNA abundance (Figure 2J), suggesting that the rate of transcription does not depend on GC content of the coding sequence found downstream of the intron. Conversely, high GC content was associated with stabilization in unspliced constructs (Figure 2K). Taken together, these experiments show that splicing preferentially increases the expression of GC-poor synonymous variants at a post-transcriptional level.

### High GC content at the 5’ end correlates with efficient expression

To further explore the sequence determinants of expression, we assembled a pool of 217 synonymous variants of GFP that included the 22 variants studied above, 137 variants from our earlier study (Kudla et al., 2009), and 58 additional variants. We cloned the collection into plasmids with and without a 5’ UTR intron. We then established pools of HeLa Flp-In T-REx cells that stably express these constructs from a single genomic locus under a doxycycline-inducible promoter and measured the protein levels of all variants by Flow-Seq (Kosuri et al., 2013). We also performed Flow-Seq in HEK293 cells using the intronless constructs only. In Flow-Seq, a pool of cells is sorted by FACS into bins of increasing fluorescence and the distribution of variants in each bin is probed by amplicon sequencing to quantify protein abundance (Figure 3A). All variants could be quantified with good technical and biological reproducibility and high correlation was found between Flow-Seq and spectrofluorometric measurement of individual constructs (Figure S4). Most variants showed the expected unimodal distribution across fluorescence bins, but some variants showed bimodal distributions, possibly indicative of gene silencing in a fraction of cells.

**Figure 3.**
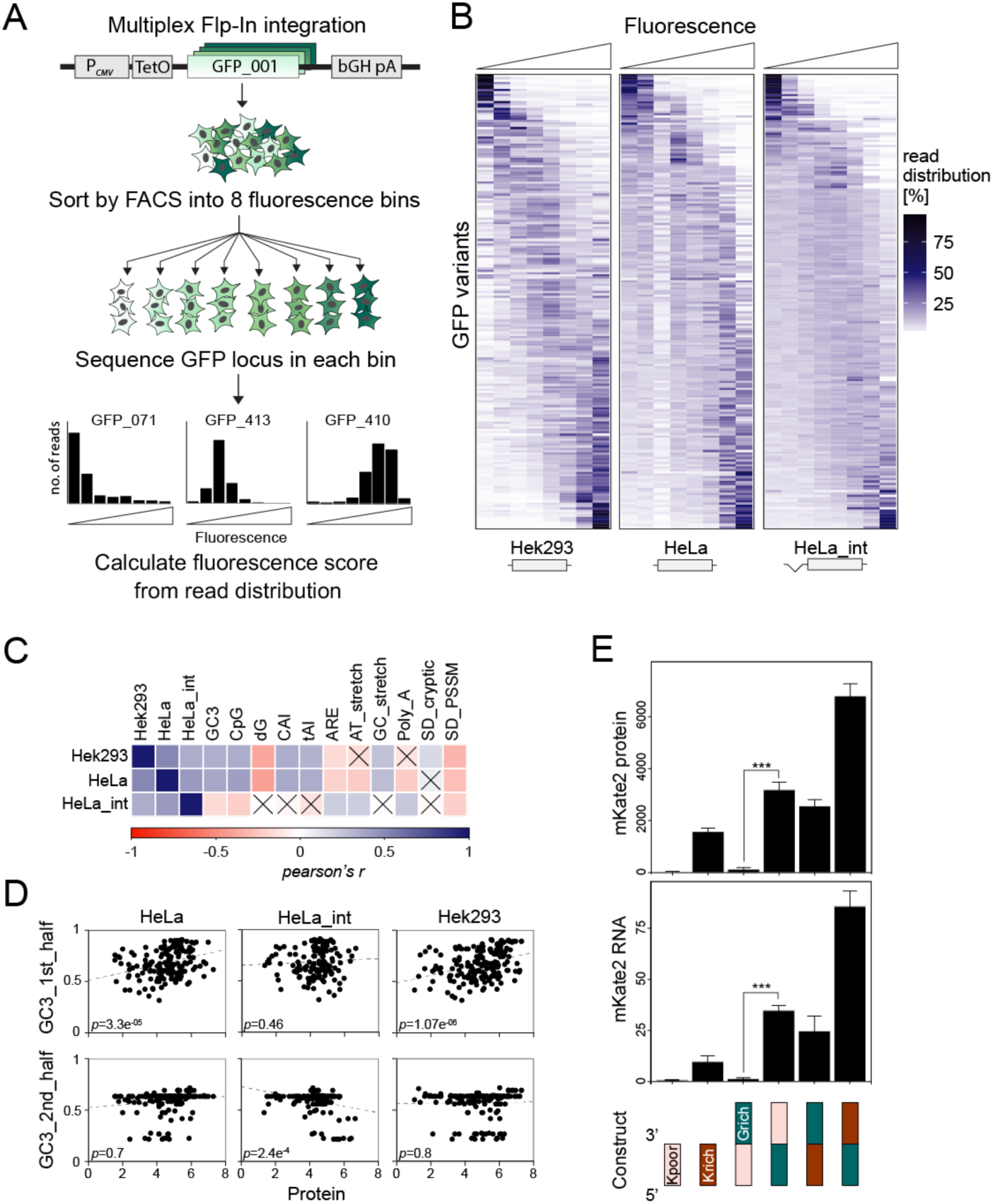
Splicing- and position-dependent effects of codon usage on protein production. (A) Schematic outline of Flow-Seq experimental workflow. Stable HeLa and HEK293 cell pools expressing 217 GFP variants were established using a multiplex Flp-In integration approach. 24h post-induction, cells are sorted by FACS into 8 fluorescence bins, genomic DNA extracted followed by high-throughput sequencing of the GFP locus. Individual fluorescence scores for each variant are calculated from normalised read distributions. (See Methods and Figure S4). (B) Heatmap representation of Flow-Seq results. Rows represent normalised read distributions of individual GFP variants across 8 fluorescence bins (columns). The average difference between lowest and highest fluorescence bin equals about 100-fold. Data shown represents the average of 3 Flow-Seq measurements for HeLa cells, the average of 3 Flow-Seq experiments for HeLa with intron and 1 experiment for HEK293 cells. (C) Pearson’s correlation matrix of experimental measurements obtained by Flow-Seq and sequence covariates. The colour of squares indicates the correlation coefficient; crosses indicate non-significant correlations. (D) Correlations between Flow-Seq measurements and GC3 content of 1^st^ (nt 1-360) and 2^nd^ (nt 361 −720) halves of GFP sequences. (E) Protein and mRNA measurements of translational fusion constructs between GC-poor (30% GC3, Kpoor) and GC-rich (85% GC3, Krich) variants of mKate2 with a GC-rich variant of GFP (97% GC3, Grich). Data represents the mean of 3 replicates + SEM (see also Figure S5).

All Flow-Seq experiments showed substantial variation of expression between synonymous variants of GFP (Figure 3B). GFP protein levels in HeLa cells (with intron), HeLa cells (without intron), and HEK293 cells (without intron) were all correlated with each other, but the moderate degree of correlation (r=0.51 HEK293 (without intron) vs HeLa (without intron); r=0.36 Hela (with intron) vs HeLa (without intron)) suggests that the effects of codon usage on expression are modulated by splicing and by cell line identity -in agreement with prior observations of tissue-specific codon usage (Burow et al., 2018; Gingold et al., 2014; Plotkin et al., 2004; Rudolph et al., 2016). Flow-Seq of unspliced variants in HeLa and HEK cells confirms the positive correlation of synonymous site GC-content with expression (Figure 3C). In contrast to the results reported by us and others in bacteria and yeast (Goodman et al., 2013; Kudla et al., 2009; Shah et al., 2013), but consistently with the positive correlation between GC content and expression, strong mRNA folding near the beginning of the coding sequence correlated with increased expression (Spearman’s ρ = 0.27 in HeLa cells; ρ = 0.4 in HEK293 cells). Expression was positively correlated with CpG content and codon adaptation index (CAI), and negatively correlated with the estimated density of ARE elements or cryptic splice sites. Because of the strong correlation between GC content, CpG content, CAI and mRNA folding energy, a multiple regression analysis could not resolve which of these properties was causally related to expression. Interestingly, for intron-containing variants, there was no correlation, or a weak negative correlation, between expression and GC content, CpG content, CAI, and mRNA folding energy (Figure 3C).

Some of the variants analysed by Flow-Seq featured large regional variation in GC content (Figure S5A) and we asked whether the localization of low-GC and high-GC regions within the coding sequence influences expression. We found that the GC3 content in the first half of the coding sequence (nt 1-360), but not in the second half (nt 361-720), was positively correlated with expression of intronless GFP variants in the HeLa and HEK293 cells (Figure 3D). The GC3 content in the first half of the gene showed no correlation with expression in the intron-containing constructs.

To further test whether the GC content at the 5’ end of genes has a particularly important effect on expression, we constructed in-frame fusions between GC-rich and GC-poor variants of GFP and mKate2 genes and quantified their protein and mRNA abundance in transient transfection experiments. Expression levels showed a striking dependence on the GC content profile. mKate2_GCpoor showed undetectable expression on its own or as a 5’-terminal fusion with GC-rich GFP, but it was efficiently expressed as a 3’-terminal fusion with GC-rich GFP (Figure 3E). By contrast, mKate2_GCrich was efficiently expressed both as 5’-terminal and 3’-terminal fusion. Analogous experiments with GC-rich and GC-poor variants of GFP fused to mKate2_GCrich led to similar conclusions (Figure S5B). The differences in protein levels between the fusion constructs could be explained by differences in mRNA abundance (Figure 3E).

### High GC content leads to cytoplasmic enrichment of mRNA and higher ribosome association

Using the pooled HeLa cells used in Flow-Seq, we then analysed the effects of GC content on mRNA localization. We separated the pools into nuclear and cytoplasmic fractions, isolated RNA and performed amplicon sequencing of each fraction to analyse mRNA localization of each GFP variant. Analysis of fractions showed the expected enrichment of the lncRNA MALAT1 in the nucleus, and of tRNA in the cytoplasm, confirming the quality of fractionations (Figure 4A). For each GFP variant, we calculated the relative cytoplasmic concentration of its mRNA (RCC) as the ratio of cytoplasmic read counts to the sum of reads from both fractions (RCC = c_cyto / (c_cyto+c_nuc); Figure 4B). A value of 0 therefore indicates 100% nuclear retention, whereas a value of 1 indicates 100% cytoplasmic localisation. In the absence of splicing, RCC scores ranged from 0.09 to 0.64 and RCC correlated significantly with GC content (r=0.51, p=3.85×10^−13^, Figure 4C). In the presence of a 5’ UTR intron, we observed a significant increase in RCC score for GFP variants with low GC content, but no increase in RCC for GC-rich variants (Figure 4D). GC3 content at the beginning of the coding sequence was significantly correlated with RCC in the absence of splicing (r=0.5, p=2.0×10^−11^), but not in the presence of splicing (R<0.01, p=0.48; Figure S5). Thus, high GC content at the 5’ end of genes increases gene expression in part through facilitating the cytoplasmic localization of mRNA.

**Figure 4.**
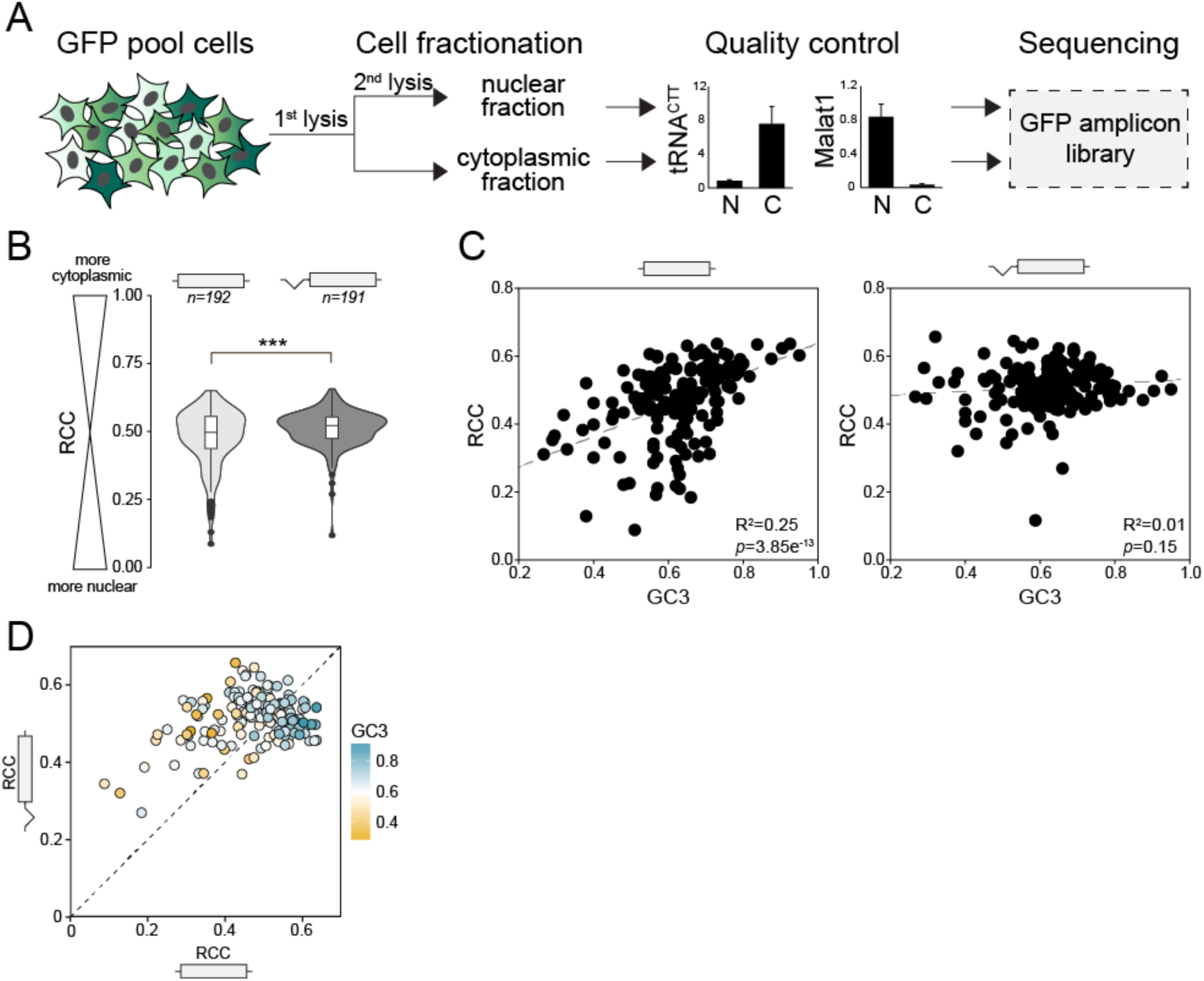
High GC content increases cytoplasmic localisation of mRNA. (A) Stable HeLa pools expressing 217 GFP variants +/-intron were fractionated into nuclear and cytoplasmic portions before RNA extraction. Specific markers of subcellular compartments were quantified by qRT-PCR before amplicon-library preparation (see also Methods). (B) Relative cytoplasmic concentration (RCC) of unspliced and spliced GFP variants. Data represents the mean of 2 replicates. ***p=2×10^−6^. (C) Correlation between GC3 content and RCC for unspliced and spliced GFP RNA. Data points represent the means of 2 replicates. (D) Correlation between RCC scores of unspliced and spliced GFP (R^2^=0.1, p=2.6×10^−5^).

To assess whether GC content also affects translational dynamics, we performed polysome profiling on HEK293 GFP pool cells using sucrose gradient fractionation (Figure 5A). qRT-PCR analyses of RNA extracted from all collected fractions showed a broad distribution of GFP across fractions, with enrichment within polysome-associated fractions. In order to determine distribution patterns of individual GFP variants, RNA from several fractions was pooled (as indicated in Figure 5B) and subjected to high-throughput sequencing. The resulting read distribution indicates that GC-rich variants are associated with denser polysomal fractions (ribosome density, Figure 5C, left panel; R^2^=0.55, p < 2.2×10^−16^) and are more likely to be translated (ribosome association, Figure 5C, right panel; R^2^=0.28, p<9.03×10^−15^), compared to GC-poor variants. This suggests that enhanced translational dynamics also contribute to more efficient expression of GC-rich genes.

**Figure 5.**
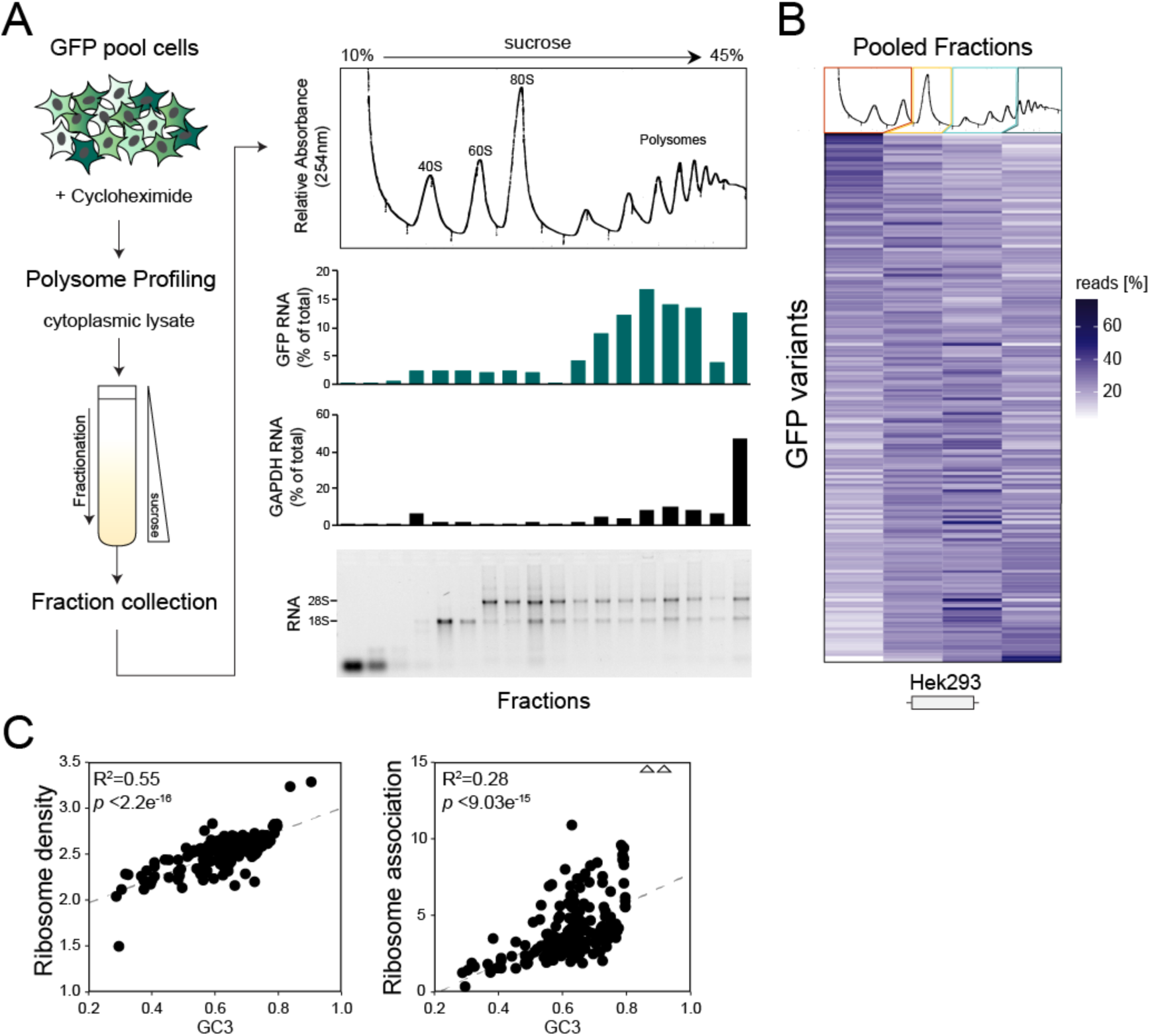
High GC content leads to increased ribosome association. (A) (Left) A stable pool of HEK293 cells expressing 217 unspliced GFP variants was subjected to polysome profiling using sucrose gradient centrifugation. (Right, from top to bottom) UV absorbance profile, GFP mRNA abundance, GAPDH mRNA abundance, ethidium bromide staining of gradient fractions. GFP and GAPDH mRNA were quantified by qRT-PCR. (B) RNA from collected fractions was combined into 4 pools (as indicated by coloured boxes) before amplicon library preparation for high-throughput sequencing: unbound ribonucleoprotein complexes (red), monosomes (yellow), light polysomes (light green) and heavy polysomes (dark green). Resulting read distributions (in %) for GFP variants are represented as heatmap. (C) Correlation plot between mean ribosome density (left panel) and ribosome association (right panel) of GFP variants and their corresponding GC3 content.

### The expression fate of endogenous RNA depends on splicing, nucleotide composition, and cell type

To test whether splicing-and position-dependent effects of codon usage can also be observed among human genes, we turned to genome-wide measurements of expression at endogenous human loci and related these measurements to codon usage and splicing. Although the correlations between GC content and expression depended on the experimental measure and type of cells under study, we find that GC4 content usually has a more positive effect on gene expression in unspliced genes relative to spliced ones (Figure 6, Table S1). In particular, unspliced mRNAs show a more positive/less negative correlation of GC4 with transcription initiation (GRO-cap data); cytoplasmic stability (exosome mutant); RNA (whole cell RNA-seq); cytoplasmic enrichment (cell fractionation), translation rate (ribosome profiling vs whole cell RNA-seq); and protein amount (mass-spec). These analyses suggest that GC4 content has an effect on the RNA abundance of intronless mRNA molecules, which is carried through to the protein expression. Taken together, these genome-wide analyses support our observation of a splicing-dependent relationship between codon usage and expression in human cells.

**Figure 6.**
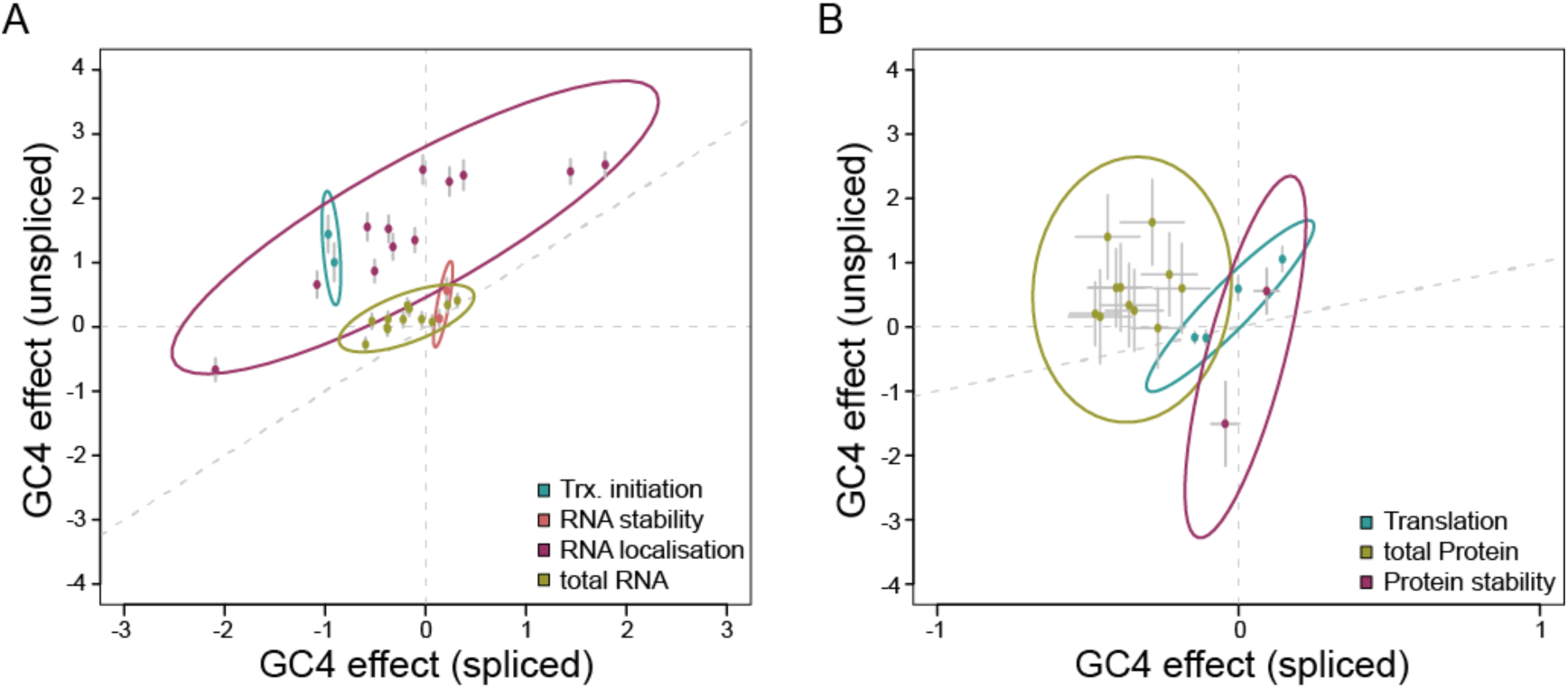
Splicing-dependent codon usage shapes global gene expression. Plots showing the effect of GC4 content on the expression of unspliced (x-axis) and spliced (y-axis) endogenous human genes, both on RNA and protein level. Each point corresponds to the regression coefficient of an individual experiment (cell line and/or biological replicate). Error bars indicate the standard error of these regression coefficients. Surrounding ellipses indicate the 95% confidence interval for 1,000 bootstraps of underlying data (see Methods, Figure S6 and Table S1). The diagonal indicates x=y.

## Discussion

We have shown that the effects of GC content on gene expression in human cells are splicing-dependent (the effect is larger in unspliced genes compared to spliced genes) and position-dependent (the effect is larger at the 5’ end of genes than at the 3’ end). In addition, human genes show striking patterns of codon usage, which differ between spliced and unspliced genes and between first and subsequent exons. Our results have implications for the understanding of the evolution of human genes and the functional consequences of synonymous codon usage.

### Mechanisms of splicing-and position-dependent effects of codon usage

Specific patterns of codon usage have previously been found at the 5’ ends of genes in bacteria, yeast and other species (Gu et al., 2010; Kudla et al., 2009; Tuller et al., 2010). In bacteria and yeast, strong mRNA folding near the start codon prevents ribosome binding and reduces translation efficiency, resulting in selection against strongly folded 5’ mRNA regions (Kudla et al., 2009; Shah et al., 2013). In addition a “ramp” of rare codons has been observed near the 5’ end of RNAs in multiple species, with a possible role in preventing a wasteful accumulation of ribosomes on mRNAs (Tuller et al., 2010) or reducing the strength of mRNA folding (Bentele et al., 2013). These phenomena cannot explain our results in human, because both the folding energy and codon ramp models predict low GC content near the start codon, whereas we observe high GC content within first exons of human protein-coding genes (Figure 1B). Furthermore, our experiments show that high GC content near the start codon increases expression, whereas the folding energy and codon ramp models would predict low expression.

We propose instead that splicing-and position-dependent effects of GC content are explained by co-transcriptional or early post-transcriptional events in the lifetime of an mRNA. Using matched reporter gene libraries, we show that most, but not all, variants show an increase in expression when spliced. Splicing typically increases the expression of AT-rich variants, but it does not further increase the expression of GC-rich transcripts, which suggests that splicing and high GC content influence expression through at least one common mechanism. Splicing increases transcription (Kwek et al., 2002), prevents nuclear degradation (Nott et al., 2003), facilitates nuclear-cytoplasmic mRNA export through the Aly/REF-TREX pathway (Muller-McNicoll et al., 2016), and stimulates translation (Nott et al., 2004). High GC content might increase RNA polymerase processivity (Bauer et al., 2010; Zhou et al., 2016); GC-rich variants are less likely to contain cryptic polyadenylation sites (consensus sequence: AAUAAA) or destabilizing AU-Rich Elements (AREs) (Higgs et al., 1983); and high GC content near the 5’ end may also facilitate cytoplasmic localisation of mRNA. GC-rich sequence elements of endogenous unspliced genes were previously shown to route transcripts into the splicing-independent ALREX nuclear export pathway, allowing efficient cytoplasmic accumulation (Palazzo et al., 2007). In agreement with this, low expression caused by inhibitory sequence features (such as low GC-content) can be rescued by extending the mRNA at the 5’end with a GC-rich sequence (Figure 3D,E, Figure S5). This may act as a compensatory mechanism when gene expression cannot rely on the positive regulatory effects of splicing (Palazzo and Akef, 2012). In contrast, it was recently shown that binding of HNRNPK to the GC-rich SIRLOIN motif leads to nuclear enrichment of lncRNAs (and also some mRNAs) (Lubelsky and Ulitsky, 2018). Our genomic analyses of lncRNA sequences do not show the same splicing-dependent compositional patterns as observed in mRNAs and it is therefore likely that antagonistic pathways act simultaneously in shaping the RNA expression landscape. Thus, we propose that the genomic patterns and their consequences on gene expression reported here are general features of protein-coding genes.

Recent studies also highlight patterns of codon usage as major determinants of RNA stability in yeast (Presnyak et al., 2015), zebrafish (Mishima and Tomari, 2016) and other species (Bazzini et al., 2016). The usage of less common, ‘non-optimal’ codons within transcripts was shown to control poly-A tail length and RNA half-life in a translation-dependent manner through the coupled activity of different CCR4-NOT nucleases (Radhakrishnan et al., 2016; Webster et al., 2018). Consistent with these findings, we observed that CAI is positively correlated with mRNA expression levels in human cells. However, it remains to be seen whether the correlation of CAI with mRNA expression depends on translation. Because of the strong correlation between GC content and CAI, it is difficult to disentangle independent contributions of these variables. Additionally, we find that the correlation between GC content (or CAI) and expression is position-and splicing-dependent, whereas no evidence for such context-dependence has been reported for the CCR4-NOT-mediated mechanism.

Other instances in which the effects of codon usage are context-dependent have been described. Most notably, tRNA populations and transcriptome codon usage patterns were shown to differ between mammalian tissues (Dittmar et al., 2006; Gingold et al., 2014; Plotkin et al., 2004; Rudolph et al., 2016). Intriguingly, genes preferentially expressed in proliferating cells and tissue-specific genes tend to be AT-rich, whereas genes expressed in differentiated cell types and housekeeping genes are more GC-rich (Gingold et al., 2014; Vinogradov, 2003). Although these differences have been interpreted in terms of the match between codon usage and cellular tRNA pools, it is plausible that translation-independent mechanisms contribute to context-dependent effects of codon usage. Accordingly, in Drosophila, codon optimality determines mRNA stability in whole cell embryos, but not in the nervous system, independent of tRNA abundance (Burow et al., 2018). Recently, it was shown that Zinc-finger Antiviral Protein (ZAP) selectively recognises high CpG-containing viral transcripts as a mechanism to distinguish self from non-self (Takata et al., 2017). We speculate that similar regulatory proteins and mechanisms exist for cellular expressed genes. The cell lines used in the present study, HeLa and HEK293, are both rapidly proliferating and experimental results are correlated (r=0.36, Flow-Seq data), but divergent expression of some GFP variants was also observed. Similarly, the effect size of GC content on the expression of endogenously expressed genes varies with cell type. It would be interesting to compare the expression of our variants in other cell types to further address the question of tissue-specific codon usage and adaptation to tRNA pools.

### Implications for the evolution of protein-coding genes

The fact that long, multi-exon genes are often found in GC-poor regions of the genome might result from regional mutation bias. However, an alternative explanation is possible: GC-poor genes may be under selective pressure to retain their introns, and intronless genes may experience selective pressure to increase their GC content. These possibilities are supported by multiple observations: Firstly, endogenous intronless genes are on average more GC-rich than intron-containing genes. Secondly, the GC content of functional (but not non-functional) retrogenes is higher compared to their respective intron-containing parental genes, which cannot be explained by a systematic integration bias. Thirdly, in genome-wide analysis, correlations between GC-content and expression are generally more positive (or less negative) for unspliced compared to spliced genes. Taken together, this suggests that for the long-term success of an unspliced gene (i.e. stable conservation of expression and functionality) an increase in GC content is essential. By contrast, splicing allows genes to remain functional even when mutation bias or other mechanisms lead to a decrease of their GC content.

## Materials and Methods

### Genes and plasmids

The library of 217 synonymous GFP variants used here consists of 138 variants from an earlier study (Kudla et al., 2009), 59 new variants assembled using the same PCR-based method as in (Kudla et al., 2009), and 22 variants that were designed *in silico* and ordered as synthetic gene fragments (gBlocks) from Integrated DNA Technologies (IDT) (Mittal et al., 2018). Each of the 22 variants was designed by setting a target GC3 content (between 25 and 95%) and randomly replacing each codon with one of its synonymous codons, such that the expected GC3 content at each codon position corresponded to the target GC3 content. For example, to design a GFP variant with GC3 content of 25%, each glycine codon was replaced with one of the four synonymous glycine codons with the following probabilities: GGA, 37.5%; GGC, 12.5%, GGG, 12.5%; GGT, 37.5%. We also generated 23 mKate2 sequences using an analogous procedure and ordered the variants as gBlocks from IDT. All the genes were cloned into the Gateway Entry vector pGK3 (Kudla et al., 2009).

### Construction of transient expression vectors

Plasmids used in transient transfection experiments are based on pCI-neo (Promega), a CMV-driven mammalian expression vector that contains a chimeric intron upstream of the multiple cloning site (MCS) within the 5’UTR. This intron consists of the 5’ splice donor site from the first intron of the human beta-globin gene and the branch and 3’ splice acceptor site from the intron of immunoglobulin gene heavy chain variable region (see pCI-neo vector technical bulletin, Promega). This vector was adapted to be compatible with Gateway recombination cloning by inserting the Gateway-destination cassette, RfA, using the unique EcoRV and SmaI restriction sites present within the MCS of pCI-neo, generating pCM2. This plasmid was then further modified by removing the intron contained within the 5’UTR by site-directed deletion mutagenesis using Phusion-Taq (ThermoScientific) and primers ‘pCI_del_F’ and ‘pCI_del_R’ (see Supplementary Table 2 for list of all primers used), generating plasmid pCM1.

To be able to normalise spectrophotometric measurements from single GFP transfection experiments, pCM1 and pCM2 were further modified to contain a separate expression cassette driving the expression of a second fluorescent reporter gene, mKate2. The mKate2 gene cassette from pmKate2-N (Evrogen) was inserted via Gibson assembly cloning: First, the entire mKate2 expression cassette was amplified using primers ‘mKate2_gibs_F’ and ‘mKate2_gibs_R’ which add overhangs homologous to the pCM insertion site. Next, pCM1 and pCM2 were linearised by PCR using primers ‘pCI_gib_F’ and ‘pCI_gib_R’. All PCR products were purified using the Qiagen PCR purification kit and fragments with homologous sites recombined using the Gibson assembly cloning kit (NEB) according to manufacturer’s instructions (NEB). Successful integration was validated by Sanger sequencing. This generated plasmids pCM3 (-intron, +mKate2) and pCM4 (+intron, +mKate2).

### Transient plasmid transfections for spectrofluorometric measurements

Plasmids for transient expression of fluorescent genes were transfected into HeLa cells grown in 96-well plates. Per plasmid construct, 3 technical replicates were tested by reverse transfection. Enough transfection mix for 4 wells was prepared by diluting 280ng plasmid DNA in 40ul OptiMem (Gibco). 1ul Lipofectamine2000 (Invitrogen; 0.25ul per well) was diluted in 40ul OptiMem and incubated for 5min at room temperature. Both plasmid and Lipofectamine2000 dilutions were then mixed (80ul total volume) and further incubated for 20-30min. 20ul of transfection complex was then pipetted into 3 wells before adding 200ul of HeLa cell suspension (45,000 cells/ml; 9,000 cells/well) in phenol red-free DMEM (Biochrom, F0475). Media was exchanged 3-4h post-transfection to reduce toxicity. Cells were then grown for a further 24h or 48h at 37C, 5% CO2.

After incubation, cells were lysed by removing media and adding 200ul of cell lysis buffer (25mM Tris, pH 7.4, 150mM NaCl, 1% Triton X-100, 1mM EDTA, pH 8). Fluorescence readings were obtained using a Tecan Infinite M200pro multimode plate reader. The plate was first incubated under gentle shaking for 15min followed by fluorescence measurements using the following settings: Ex486nm/Em 515nm for GFP and Ex588nm/Em633nm for mKate2; reading mode: bottom; number of reads: 10 per well; gain: optimal.

For data analysis, measurements of untransfected cells were subtracted as background from all other wells. For comparability of different plates within a set of experiments, the same 3 genes were transfected on every plate to account for technical variability. In the screen of individual GFP variants (see Figure 2), GFP measurements were divided by mKate2 measurements from same wells to reduce noise caused by well-to-well variation in transfection efficiency, but similar results were obtained without normalisation.

### Transient transfections and RNA extraction for qRT-PCR analysis

HeLa cells were reverse transfected in 12-well plates using 800ng plasmid DNA and 2ul Lipofectamine 2000 (Invitrogen). DNA and Lipofectamine 2000 were diluted in 100ul OptiMEM (Gibco) each, incubated for 5min, mixed and further incubated for 20min. The transfection complex was then added to each well before adding 10^5^ HeLa cells. Cells were incubated for 24h at 37C, 5% CO2 before harvesting. Cells were then harvested by adding 1ml Trizol reagent (Life technologies). RNA was extracted according to manufacturer’s instructions. Resulting RNA was further treated with DNAse I using the Turbo DNase kit (Ambion) to remove any residual plasmid and genomic DNA.

### qRT-PCR analysis

cDNA for qRT-PCR analysis was prepared using SuperScript III Reverse Transcriptase (Life technologies) according to the manufacturer’s recommendations with 500ng total RNA as template and 500ng random hexamers (Promega). All qRT-PCRs were carried out on a Roche LightCycler 480 using Roche LightCycler480 SYBR Green I Master Mix and 0.3uM gene-specific primers. Samples were analysed in triplicate as 20ul reactions, using 2ul of diluted cDNA. Cycling settings: DNA was first denatured for 5min at 95°C before entering a cycle (50-60x) of denaturing for 10sec at 95°C, annealing for 7sec at 55-60°C (depending on primers used), extension for 10sec at 72°C and data acquisition. DNA was then gradually heated up by 2.20 °C/s from 65 to 95°C for 5sec each and data continuously collected (Melting curve analysis). Data was evaluated using the comparative Ct method (Livak and Schmittgen, 2001). RNA measurements from transient transfection experiments were normalised to the abundance of neomycin RNA, which is expressed from the same plasmid, to control for differences in transfection efficiency (primers ‘Neo_F’ and ‘Neo_R’).

### Subcellular fractionation

This protocol is based on the cellular fractionation protocol published by (Gagnon et al., 2014) but includes a further clean-up step using a sucrose cushion as described by (Zaghlool et al., 2013) and a second lysis step as described by (Wang et al., 2006). Cell lysis and nuclear integrity was monitored throughout by light microscopy following Trypan blue staining (Sigma). Cells were grown in 10cm plates for 24h to about 90% confluency. Cells were then washed with PBS and trypsinised briefly using 1ml of 1xTrypsin/EDTA. After stopping the reaction with 5ml DMEM, cells were transferred into 15ml falcon tubes and collected by spinning at 100g for 5min. Resulting cell pellets were resuspended in 500ul ice-cold PBS, transferred into 1.5ml reaction tubes and spun at 500g for 5min, 4°C. The supernatant was discarded and cells resuspended in 250ul HLB (10mM Tris (pH 7.5), 10mM NaCl, 3mM MgCl2, 0.5% (v/v) NP40, 10% (v/v) Glycerol, 0.32M sucrose) containing 10% RNase inhibitors (RNasin Plus, Life Tech) by gently vortexing. Samples were then incubated on ice for 10min. After incubation, samples were vortexed gently, spun at 1000g for 3min, 4°C, and supernatants and pellets were processed separately as indicated in a) and b) below.

a) Cytoplasmic extract:

The supernatant was carefully layered over 250ul of a 1.6M sucrose cushion and spun at 21,000g for 5min. The supernatant was then transferred into a fresh 1.5ml tube and 1ml Trizol was added and mixed by vortexing.

b) Nuclear extract:

The pellets were washed 3 times with HLB containing RNase inhibitors by gently pipetting up and down 10 times followed by a spin at 300g for 2min. After the 3rd wash, nuclei were resuspended in 250ul HLB and 25ul (10%) of detergent mix (3.3% (wt/wt) sodium deoxycholate/6.6% (vol/vol) Tween 40) dropwise added while vortexing slowly (600rpm). Nuclei were then incubated for 5min on ice before spinning at 500g for 2min. The supernatant was discarded and pellets resuspended in 1ml Trizol (Ambion) by vortexing. 10ul 0.5M EDTA are added to each nuclear sample in Trizol and tubes heated to 65°C for 10min to disrupt very strong Protein-RNA and DNA-RNA interactions. Tubes were then left to reach room temperature and RNA was extracted following the manufacturer’s instructions.

### Transcription inhibition assay

HeLa T-Rex Flp-in cell lines were grown to 80-90% confluency in 6 well for 24h before treatment with 500nM Triptolide (Sigma). Cells were harvested at indicated time points and RNA extracted using Trizol reagent (Ambion). Control cells were treated with the equal volume of DMSO (drug carrier). To assess transcript levels, qRT-PCR was performed as described above. GFP levels were normalised to levels of 7SK, a RNA polymerase III-transcribed non-coding RNA, whose expression levels are not affected by Triptolide treatment. Relative transcript levels of c-Myc are shown as an example of a relatively unstable transcript.

### Generation of stable Flp-in cell lines

We adopted a multiplex-Gateway integration method to create a pool of 217 GFP plasmids which are compatible with the T-Rex Flp-in system (Invitrogen) for creating stable, doxycycline-inducible cell lines, in which each variant is expressed from the same genomic locus, allowing direct comparison of expression levels.

pcDNA5/FRT/TO/DEST (Aleksandra Helwak, University of Edinburgh) contains the Gateway-compatible attB destination cassette to allow the subcloning of genes from any Gateway-entry vectors. This plasmid was further modified to contain the same 5’UTR intron sequence as in pCM4 used in transient expression experiments using Gibson Assembly (NEB): the intronic sequence was amplified from pCM4 by PCR using primers ‘Gib_intr_F’ and ‘Gib_intr_R’ using Q5 High-Fidelity Polymerase (NEB). The primers added 15nt overhangs which are homologous to the ends of pcDNA5/FRT/TO/DEST when linearised with AflII. The Gibson assembly reaction was performed as per manufacturer’s instructions (NEB), generating pcDNA5/FRT/TO/DEST/INT.

217 individual GFP variants stored in Gateway-entry vector pGK3 were mixed with a concentration of 0.06ng of each GFP variant. For each pcDNA5 destination vector, a separate Gateway LR reaction was set-up in a total volume of 45ul using 500ng destination vector, 5ul LR Clonase enzyme mix, 38ul of the mixed 217 pGK3-GFP plasmids and TE (pH 8). The reactions were incubated at 25C overnight followed by Proteinase K digest (5ul, LR Clonase kit) for 10min at 37C. The total 50ul reaction mix was transformed into 2.5ml highly competent DH5alpha in a 15ml Falcon tube by heat-shocking cells for 2min 30s at 42C, followed by cooling on ice for 3min, before adding 10ml SOC medium and incubating while shaking for 1h at 37C. After incubation, cells were spun down at 3000g for 3min and resulting bacterial pellets resuspended in 1ml fresh SOC. 10×100ul were plated onto L-Ampicillin agar plates and incubated overnight at 37C resulting in >800 colonies per plate. Bacterial colonies were scraped off the plates and collected in a falcon tube. Plasmid DNA was extracted using a Qiagen Midiprep kit according to the manufacturer’s instructions, resulting in two plasmid pools: pCDNA5/GFPpool and pcDNA5/INT/GFPpool. Both pools were subjected to high-throughput sequencing to confirm the presence of different GFP variants.

HeLa T-Rex Flp-in cells (gifted by the Andrew Jackson lab, The University of Edinburgh) and Hek293 T-Rex Flp-in (Thermo Scientific) were grown to 80% confluency in 6 well plates. For GFP plasmid pool transfections, pCDNA5/GFPpool or pCDNA5/INT/GFPpool were mixed in a 9:1 ratio with the Flp-recombinase expression plasmid pOG44 (Invitrogen) to give 2ug in total (1.8ug pOG44 + 0.2ug pCDNA5) and diluted in OptiMEM (Gibco) to 100ul. Transfections were performed with 9ul Lipofectamine2000 (Invitrogen) and 91ul OptiMEM per well by incubating 5min at room temperature before mixing with plasmid DNA and a further 15min incubation. The transfection mix was then added dropwise to the cells. Media were replaced with conditioned media 4h post-transfection. Cells were incubated for further 48h before chemical selection to select for successful gene integration using 10ng/ul Blasticidin S (ThermoFisher) and 400mg/ml (HeLa T-Rex Flp-in) or 100mg/ml (Hek293 T-Rex Flp-in) Hygromycin B (Life Technologies). Successful selection was determined by monitoring cell death in untransfected cells. Chemically resistant cells represent pools of cell lines expressing different GFP variants from the same genomic locus. High-throughput sequencing of the GFP integration site within each generated cell line pool confirmed the successful integration of all variants.

HeLa T-Rex Flp-in and Hek293 T-Rex Flp-in cell lines expressing two individual GFP variants (GC3=96% and GC3=36%; see Supplementary Figure 3) as spliced and unspliced transcripts were generated using the same protocol.

### Flow-Seq: FACS sorting and genomic DNA extraction

80×15cm cell culture plates of HeLa T-Rex Flp-in GFP pool cells and 40×15cm cell culture plates of Hek293 T-Rex Flp-in GFP pool cells were induced with 1ug/ml Doxycyline (Sigma, D9891) in phenol red-free DMEM (Biochrom, F0475) supplemented with 10% FCS (Sigma, F-7524) and 2mM L-Glutamine. After 24h or 48h, cells were harvested by gentle trypsinisation and cells were sorted into 8 fluorescence bins using a BD FACS Aria II cell sorter. To define the range of GFP positive signal, cells without stable GFP expression were used as negative control. 80% of HeLa and 90% Hek293 GFP pool cells fell into the GFP-positive range. Each fluorescence bin was chosen to comprise roughly 10% of the GFP-positive population. The bin spacing was kept the same for the sorting of HeLa cell pools expressing unspliced and spliced GFP variants to allow direct comparisons of the fluorescence profiles of individual variants.

About 10^7^ cells per bin were collected in Polypropylene collection tubes (Falcon) coated with 1% BSA/PBS, cushioned with 200ul 20%FBS/PBS. Cell suspensions were decanted into 15ml tubes and cells collected by spinning 5min at 500g. The supernatant was transferred into fresh 15ml tubes and precipitated using 2 volumes of 100% EtOH/0.1 volume Sodium Acetate (pH 5.3) and 10ul Glycoblue (Ambion). Tubes were shaken vigorously for 10s before incubating at −20C for 15min, followed by spinning at 3000g for 20min. Resulting pellets were air-dried, resuspended in 1ml digest buffer (100mM Tris pH 8.5, 5mM EDTA, 0.2% SDS, 200mM NaCl) and then combined with the respective cell pellet. 10ul RNAse A (Qiagen, 70U) was added and samples gently rotated at 37C. After 1h, 1ul/ml Proteinase K (20mg/ml, Roche) was added to the samples before rotating a further 2h at 55C. Genomic DNA was purified 3 times by using 1 volume Phenol:Chloroform:Isoamyl alcohol (PCI, 25:24:1, Sigma). After each addition of PCI, samples were shaken vigorously for 10s before spinning at 3000g for 20min (first extraction) or 5min (all following). The resulting bottom layers including the interphase were removed before each PCI addition. After the last PCI extraction, the upper layer was transferred into a fresh 15ml tube and 1 extraction performed using 1 volume chloroform:isoamyl alcohol (CI,24:1, Sigma). After a 5min spin at 3000g, the upper layer was transferred into a fresh 15ml tube and DNA precipitated using EtOH/Sodium Acetate as before. After a 5min incubation on ice, DNA was collected by spinning for 30min at 3000g. The resulting DNA pellets were washed 2 times with 75% EtOH before air-drying and resuspending in 200ul Tris-EDTA (10mM). The quality of the extracted genomic DNA was assessed on a 0.8% Agarose/TBE gel.

### Polysome profiling

Hek293 Flp-in GFP pool cell lines were grown to 90% confluency on 15cm dishes. Cells were treated for 20min with 100ug/ul Cycloheximide before harvesting cells by removing media, washing with 2x ice-cold PBS followed by scraping cells into 1ml PBS and transferring into 1.5ml tubes. Cells were pelleted at 7000rpm, 4°C for 1min and resulting cell pellet carefully resuspended by pipetting up and down in 250ul RSB (10x RSB: 200mM Tris (pH 7.5), 1M KCl, 100mM MgCl2) containing 1/40 RNasin (40U/ul, Promega), until no clumps were visible. 250ul of polysome extraction buffer was then added (1ml 10x RSB + 50ul NP-40 (Sigma) + 9ml H2O + 1 complete mini EDTA-free protease inhibitor pill (Roche)) and lysate passed 5x through a 25G needle avoiding bubble formation. The lysate was then incubated on ice for 10min before spinning 10min at 10,000g, 4°C. The supernatant was then transferred into a fresh 1.5ml tube and the RNA concentration estimated by measuring the OD at 260nm. Sucrose gradients (10–45%) containing 20 mM Tris, pH 7.5, 10 mM MgCl2, and 100 mM KCl were made using the BioComp gradient master. 100ug of Lysate were loaded on sucrose gradients and spun at 41,000rpm for 2.5h in a Sorvall centrifuge with a SW41Ti rotor. Following centrifugation, gradients were fractionated using a BioComp gradient station model 153 (BioComp 23 Instruments, New Brunswick, Canada) by measuring cytosolic RNA at 254 nm and collecting 18 fractions.

RNA from all fractions was precipitated using 1 volume of 100% EtOH and 1ul Glycoblue (Ambion), before extracting RNA using the Trizol method (Life Technologies). Equal volumes of RNA of each fraction was run on a 1.3% Agarose/TBE gel to assess the quality of fractionation and RNA integrity. Additionally, equal volumes of RNA of each fraction were used in cDNA synthesis using SuperScript III (ThermoFisher) and 2uM gene-specific primers for GFP (‘pcDNA5-UTR_R’) and GAPDH (‘GAPDH_R’) followed by qRT-PCR analysis. For high-throughput sequencing, total RNA from collected fractions was combined in equal volumes into 4 pools (as indicated in Figure 5B; free ribonucleoprotein (RNP) complexes, monosomes, light polysomes (2-4) and heavy polysomes (5+)) before amplicon library preparation (as described below).

### High-throughput library preparation and sequencing

Sequencing libraries were generated by PCR using primers specific for GFP amplification (Supplementary Table 2) which carry the required adaptor sequences for paired-end MiSeq sequencing, as well as 6nt indices for library multiplexing. Between 6-10ug of total genomic DNA were used in multiple PCR reactions (200ng per 50ul reaction). All PCRs were performed using Accuprime Pfx (NEB) according to manufacturer’s recommendations using 0.4ul Accuprime Pfx Polymerase and 0.3uM of each primer (‘PE_PCR_left’ and ‘S_indexX_right_PEPCR’). The cycling conditions were as follows: Initial denaturation at 95C for 2min, followed by 30 cycles of denaturation at 95C for 15sec, annealing at 51C for 30sec, extension at 68C for 1min. The final extension was performed at 68C for 2min. After PCR, all reactions of the same template were pooled and 1/3 of the reaction purified using the Qiagen PCR purification kit according to the manufacturer’s instructions. DNA was eluted in 50ul H2O. Library size selection was performed using the Invitrogen E-gel system (Clonewell gels, 0.8% agarose) followed by Qiagen MinElute PCR purification. Correct fragment sizes were confirmed and quantified using the Agilent Bioanalyzer 2100 system.

For library preparation of RNA samples, 500ng RNA was first converted into cDNA using 2nmol GFP-specific primers (‘S_indexX_right_PEPCR’) using SuperScript III (Life technologies) according to manufacturer’s protocol, using 50C as extension temperature. Resulting cDNA was then treated with 1ul RNaseH (NEB) for 20min at 37C, followed by heat inactivation at 65C for 5min. Samples were diluted 1:2.5 before using 2ul as template in PCR for library preparation. A minimum of 8×50ul PCR reactions were set up and pooled for each sample before PCR purification, followed by E-gel purification as described above.

High-throughput sequencing was conducted by Edinburgh Genomics (The University of Edinburgh) and Imperial BRC Genomics facility (Imperial College London) using the Illumina MiSeq platform (2×300nt paired-end reads).

### Analysis of GFP pool experiments

Raw sequencing files (fastq files) were demultiplexed by 6nt indices by the respective sequencing facility. To remove the plasmid sequence, the second reads from paired-end sequencing were trimmed using flexbar (-as ATGTGCAGGGCCGCGAATTCTTA -ao 4 -m 15 -u 30). Reads were then mapped to the GFP library using bowtie2 (-X 750) and filtered using samtools (-f 99).

For Flow-seq data, only variants with a minimum of 1000 reads across all 8 sequencing bins were used for further analysis. For each GFP variant, the number of reads in each bin (n(i)) was multiplied by the respective bin index (i) before taking the sum and dividing by the total number of reads across all bins: Fluorescence 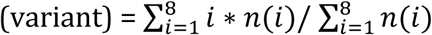

For cell fractionation experiments, only data with a minimum of 1000 reads across both cytoplasmic and nuclear fractions was used to calculate the relative cytoplasmic concentration (‘RCC’) for each variant: 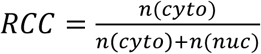

For polysome profiling, only variants with a minimum of 1000 reads across all 4 sequencing bins were used for further analysis. To estimate ribosome density, for each GFP variant, the number of reads in each bin (n(i)) was multiplied by the respective polysomal fraction index (i) before taking the sum and dividing by the total sum of reads across all fractions:

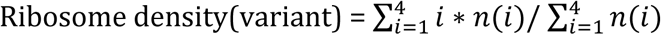

Ribosome association for each variant was calculated as the sum of reads (n) in light polysomes, heavy polysomes and monosomal fractions, divided by the sum of reads found in the free RNP fraction:

Ribosome association(variant) = (n(monosomes) + n(light polysomes) + n(heavy polysomes)) / n(free RNPs)

### Definition of calculated sequence features

GC3: GC content in the third position of codons

CpG: number of CpG dinucleotides

dG: The minimum free energy of predicted mRNA secondary structure around the start codon was calculated using the hybrid-ss-min program version 3.8 (default settings: NA = RNA, t = 37, [Na+] = 1, [Mg++] = 0, maxloop = 30, prefilter = 2/2) in the 42-nt window (−4 to 38) as in (Kudla et al., 2009).

CAI: Codon Adaptation Index (*H. sapiens*) (Sharp and Li, 1987a) was calculated using a reference list of highly expressed human genes collected from the EMBL-EBI expression atlas https://www.ebi.ac.uk/gxa.

tAI: tRNA adaptation index (dos Reis et al., 2004)

ARE: top score of ATTTA motif match in each sequence.

AT-stretch: number of times motif (AT){9} was identified in each sequence. GC-stretch: number of times motif (GC){9} was identified in each sequence.

Poly_A: number of times the position-specific scoring matrix ((47,3,0,50)(18,6,9,67)(53,12,12,23)(59,6,0,35)(70,6,6,18)) was identified in each sequence.

SD_cryptic: number of times RSGTNNHT motif was identified in each sequence. SD_PSSM: number of times the position-specific scoring matrix ((60,13,13,14)(9,3,80,7)(0,0,100,0)(0,0,0,100)(53,3,42,3)(71,8,12,9)(7,6,81,6)(6,17,21,46)) was identified in each sequence.

FIMO (http://meme-suite.org) was calculated to identify and count sequence motifs. Open-source packages available for R were used for generating correlation matrices (corrplot), heatmaps (ggplot2), boxplots (graphics/ggplot2), The GC3 of all human coding sequences (assembly: GRCg38_hg38; only CDS exons) was calculated using R package ‘seqinr’.

### Computation methods for analysis of endogenous gene expression Data Collection – see also Supplementary Table 1

1. GC4 content was calculated for each protein-coding transcript annotated in GENCODE version 19 as the GC content of the third codon position across all fourfold-degenerate codons (CT*, GT*, TC*, CC*, AC*, GC*, GA*, CC*, GC*). The core promoter of each transcript is further defined as −300 bp/+100 bp around the annotated TSS.
2. The level of transcription initiation was quantified in K562 and Gm12878 cells as the number of GRO-cap reads from the same strand which overlap the core promoter.
3. Nuclear stability was assessed using CAGE data obtained in triplicate from Egfp, Mtr4 and Rrp40 knockdowns (GSE62047; (Andersson et al., 2014)). Similarly to the approach used for the GRO-cap data, we calculated the RPKM across core promoters for each library separately. The baseMean expression for each treatment was quantified using DESeq2, where promoters with no reads across any replicate were first removed from each comparison. Nuclear stability was then assessed as the fold-change between the Egfp and Mtr4 knockdown and cytoplasmic stability by the estimated fold-change between the Mtr4 and Rrp40 knockdowns.
4. The level of the mature mRNA was quantified using RNA-seq libraries from whole cell samples (prepared as described elsewhere for HEK293 cells and downloaded from http://hgdownload.cse.ucsc.edu/goldenPath/hg19/encodeDCC/wgEncodeCshlLongRnaSeq for Gm12878, HepG2, HeLa, Huvec and K562 cells). Reads were pseudoaligned against GENCODE transcript models using Kallisto, set with 100 bootstraps. All other parameters were left at their default. Transcript expressions were extracted as the estimated TPM (tags per million) values.
5. The level of the mature mRNA in the nuclear and cytoplasmic fractions was quantified using Kallisto as previously. As transcript stability was similar in both fractions (linear regression coefficient 0.97, p < 2.2×10^−16^), nuclear export was determined as the fraction TPM from these two compartments which was present in the nuclear fraction.
6. Ribosome-sequencing data from HEK293 (GSE94460) and HeLa (GSE79664) cells were used to quantify the level of mRNA translation in these two cells. Both of these measures were determined at the gene level, and so these observations were applied to all GENCODE transcripts annotated to these associated genes. These data were normalised to the mean mRNA expression in the relevant cell types (from step 4).
7. Protein expression was assessed using mass-spectrometry data (Geiger et al., 2012) (Supp. Table 2) as the mean LFQ intensity across three replicates for each uniprot-annotated gene in each cell line for which data were available. Only data from genes where the UniProt ID is uniquely linked to a single transcript were considered in the analyses presented here.
8. Protein stability was calculated as the level of the mature protein in HEK293 and HeLa cells (step 7) relative to the mean rate of mRNA translation in these cells (step 6).

### Regression modelling

A pseudocount of 0.0001 was added to each measurement of gene expression and, excluding the nuclear export data, these values were then log2-transformed to generate a normal distribution of expression for subsequent analysis. Transcripts with an expression value of 0 were removed from downstream analysis and the resulting distributions used for regression analysis are displayed in Supplementary Figure 6. Transcripts were separated into unspliced and spliced, where splicing was defined as containing more than one exon in the GENCODE transcript model. Expression measurements were then linearly regressed against the GC4 content separately for each class of transcript and the coefficients along with their associated standard errors. These data were then bootstrapped by sampling with replacement and recalculating the regression coefficients for spliced and unspliced transcripts. The 95% confidence interval of these coefficients (discounting the standard error in these estimations) obtained by 1,000 samplings of this type was used to draw the ellipses shown in Figure 6.

### Analysis of GC content variation in the human genome

The GRCh38 sequence of the human genome, as well as the corresponding gene annotations (Ensembl release 85), was retrieved from the Ensembl FTP site (Zerbino et al., 2018). The full coding sequences (CDSs) of protein-coding genes were extracted, filtered for quality and clustered into putative paralogous families (see (Savisaar and Hurst, 2016) for full details). For all analyses, a random member was picked from each putative paralogous cluster. In addition, only one transcript isoform (the longest) was considered from each gene. Note that exon rank was always counted from the first exon of the gene, even if it was not coding. For Figure 1C, GC4 was averaged across all sites that were at the same nucleotide distance to the TSS and within an exon of the same rank. For the functional retrocopies analysis, the parent-retrocopy genes derived in (Parmley et al., 2007) were used. Pseudogenic retrocopies were retrieved from RetrogeneDB (Rosikiewicz et al., 2017). Retrocopy annotations were filtered to only leave human genes with a one-to-one ortholog in *Macaca mulatta*. Next, only ortholog pairs where both the human and the macaque copy were annotated as not having an intact reading frame and where the human copy was annotated as *KNOWN_PSEUDOGENE* were retained. For the analyses reported in Supplementary Figure 1, the functional retrocopies were also retrieved from RetrogeneDB, as we could not access genomic locations for the (Parmley et al., 2007) set. The functional retrogenes were retrieved similarly to pseudogenes, except that both the human and the macaque copy were required to have an intact open reading frame and the human copy could not be annotated as *KNOWN_PSEUDOGENE*.

Python 3.4.2. was used for data processing and R 3.1.2 was used for statistics and plotting (R Development Core Team, 2005).

## Acknowledgments

We thank Elisabeth Freyer for help with cell sorting; Andrew Jackson, Nick Gilbert and Aleksandra Helwak for gifts of cell lines and plasmids; James Brindle for technical assistance; Michael Liss and members of Kudla and Hurst groups for discussions; Edinburgh Genomics (University of Edinburgh) and the Imperial BRC Genomics facility for next-generation sequencing; and Institute of Genetics and Molecular Medicine technical support for help with media preparation and sequencing. This work was supported by the Wellcome Trust (Fellowships 097383 and 207507 to G.K.), the European Research Council (Advanced grant ERC-2014-ADG 669207 to L.D.H.), and the Medical Research Council (Grants MC_UU_00007/11 to M.S.T. and MC_UU_00007/12 to G.K. and PhD studentship to C.M.).

---

**Supplementary Figure 1.**
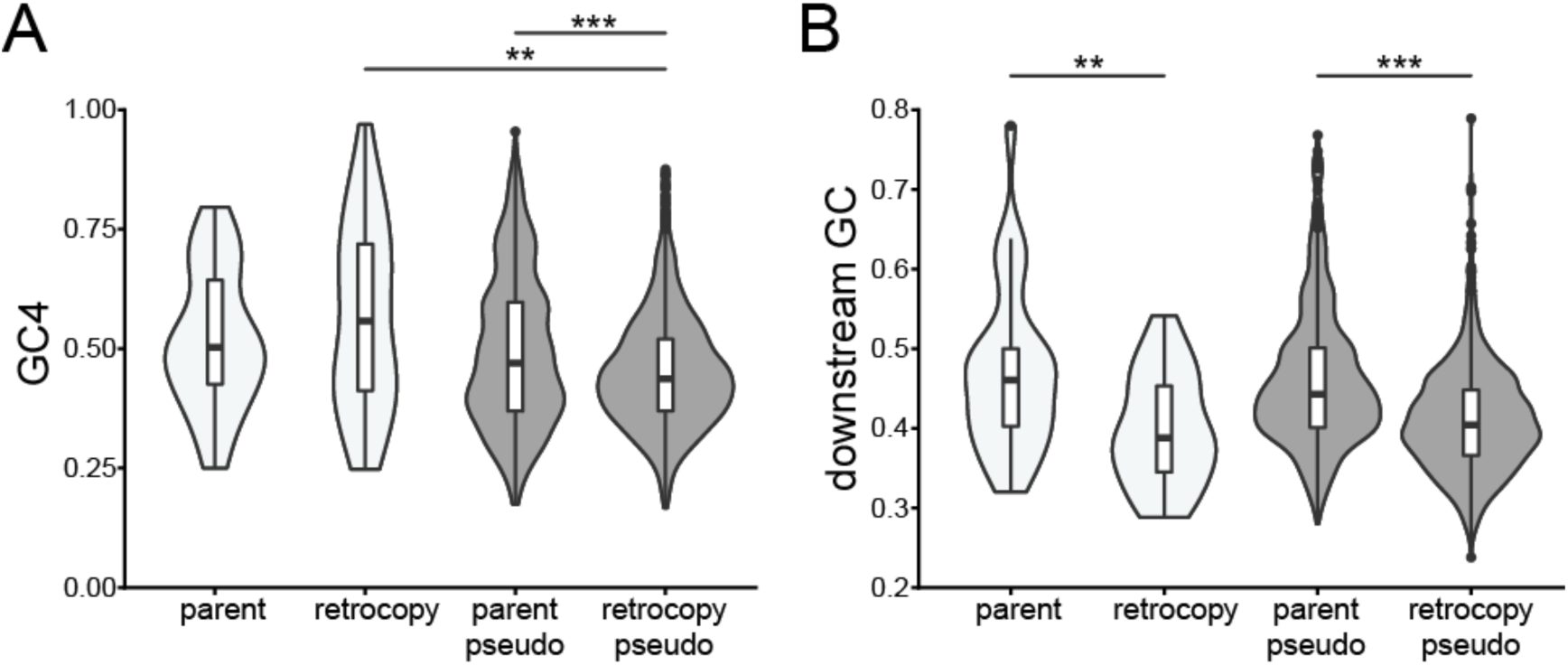
GC4 variation amongst parent-retrogene pairs and their downstream sequence. (A) GC4 content distribution across parent and retrogene pairs conserved between human and macaque. White violins indicate pairs for which retrocopies are classed as functional (p=0.26, n=31, two-tailed Wilcoxon signed-rank test), whereas grey violins correspond to pairs in which the retrocopy is classed as non-functional pseudogene (p < 2.2×10^−16.^, n=1562, two-tailed Wilcoxon signed-rank test). Note that a different retrogene dataset was used than in the main text (see Methods for details). For the human-macaque set, the difference in GC4 between parents and functional copies is in the expected direction but not significant. (B) Violin plot showing GC content within a window between 2000 and 3000nt downstream from the stop codons of functional (white, p=9.27×10^−4^, n=31, two-tailed Wilcoxon signed-rank test) and non-functional (grey, p<2.2×10^−16^, n=1562, two-tailed Wilcoxon signed-rank test) parent-retrogene pairs conserved between human and macaque.

**Supplementary Figure 2.**
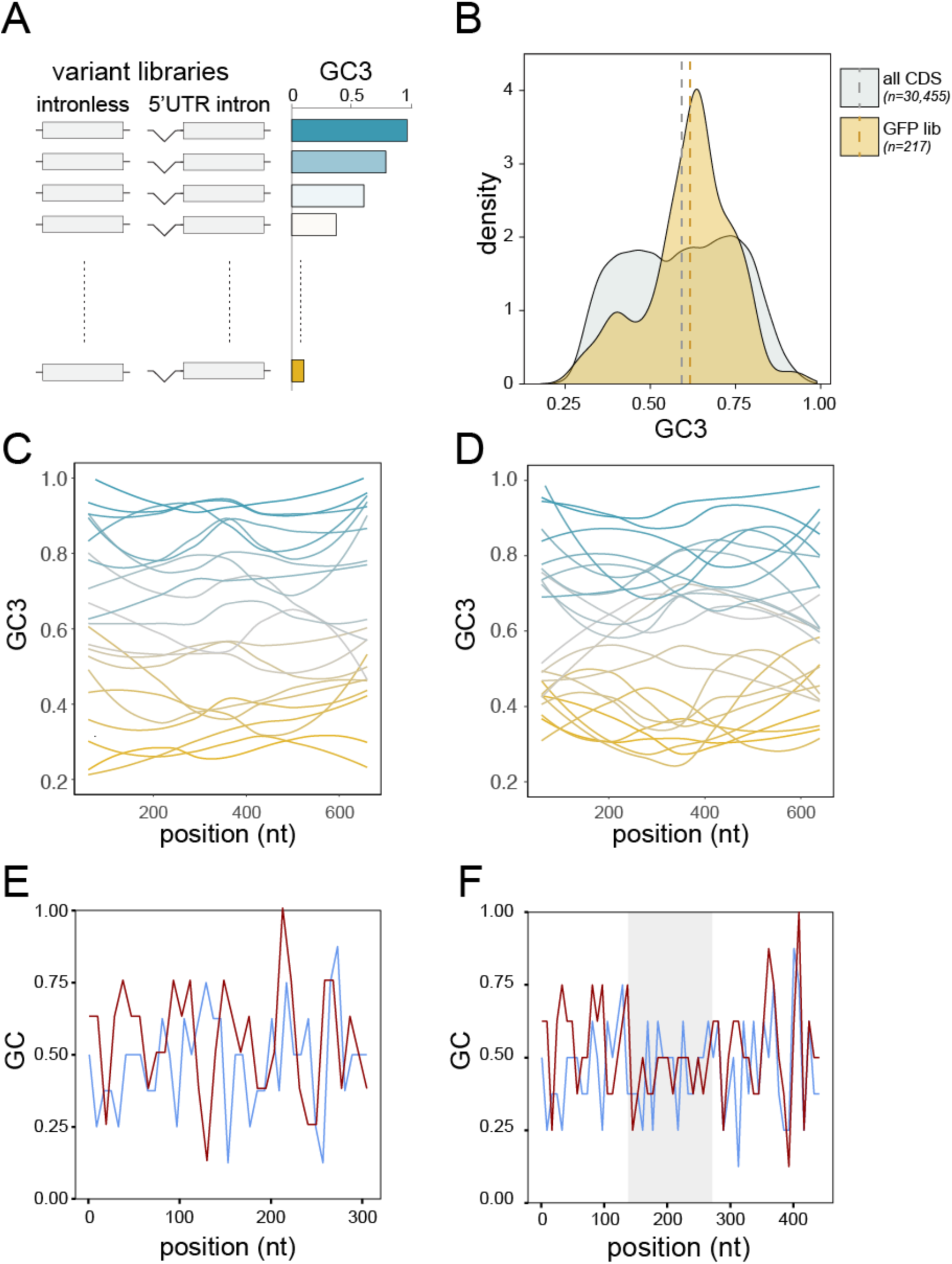
GC content variation amongst endogenous genes and reporter libraries. (A) Libraries of reporter genes with random synonymous codon usage were designed to cover a broad range of GC3 content variation. Variants were expressed with and without a synthetic 5’UTR intron. (B) GC3 content distribution amongst human consensus coding sequences (CDS; grey) in comparison to the GFP variant library used in this study (GFP lib; orange). Dashed lines indicate the mean GC3 for each data set. (C-D) Loess-smoothed GC3 profiles along the 22 GFP variants (C) and 23 mKate variants (D) that were analysed by spectrofluorometry (Figure 2). (E) Sliding window analysis of GC content in 5’UTRs of intronless expression cassettes utilised in this study. Blue: pCM3 (transient transfection, no intron); red: pcDNA5/FRT/TO/DEST (stable transfection, no intron). (F) As above, intron-containing expression cassettes. Blue: pCM4 (transient transfection, with intron); red: pcDNA5/FRT/TO/DEST/INT (stable transfection, with intron). Grey shading indicates the position of the synthetic intron.

**Supplementary Figure 3.**
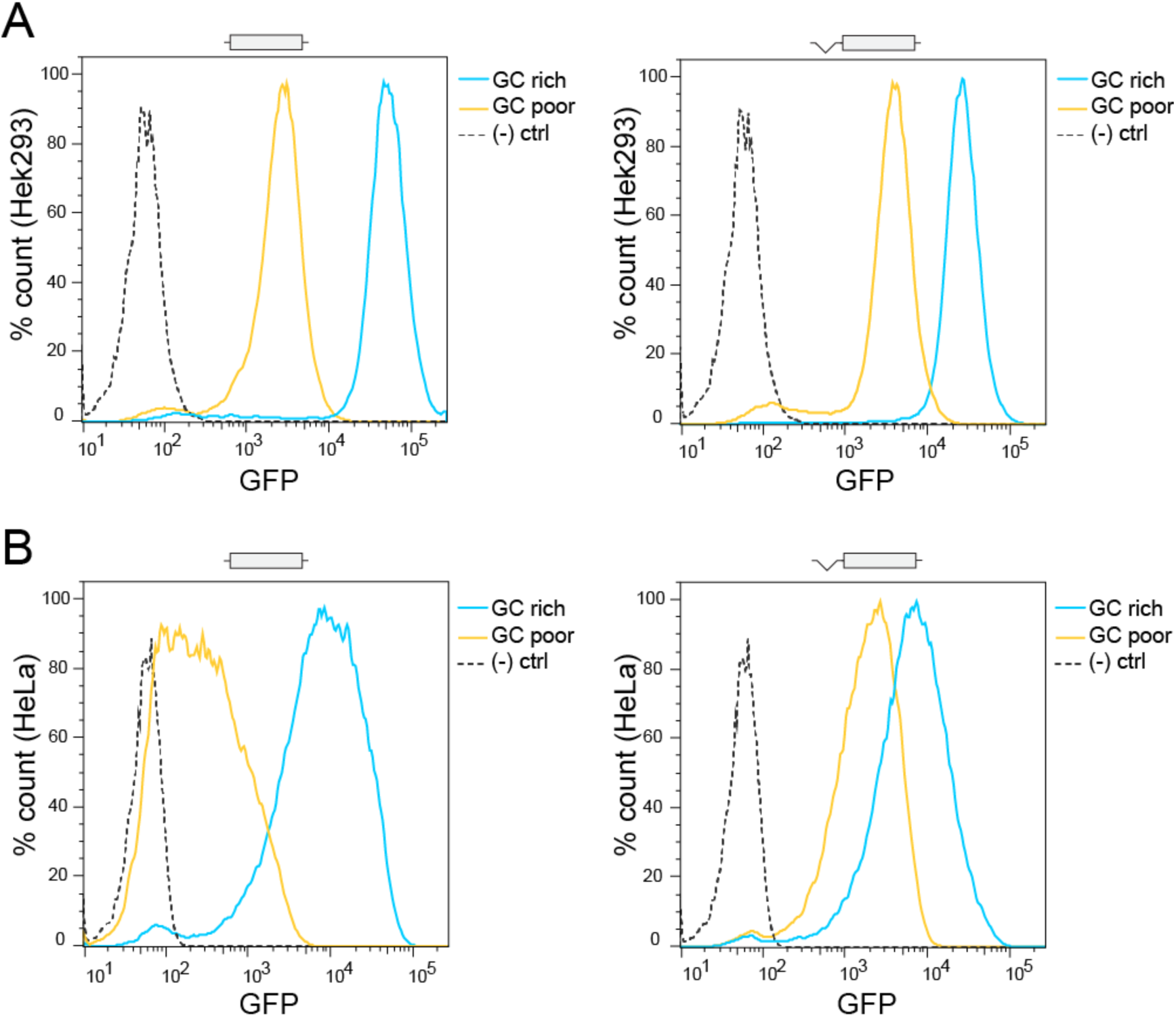
Effect of GC content on expression of fluorescent reporter genes in stably transfected cell lines. Flow cytometry measurements of two GFP variants in stably transfected HEK293 Flp-in (A) and HeLa Flp-in (B). GC poor = 33% GC3; GC rich = 97% GC3; (-)ctrl = untransfected cells. Data shows representative results of at least 3 experiments.

**Supplementary Figure 4.**
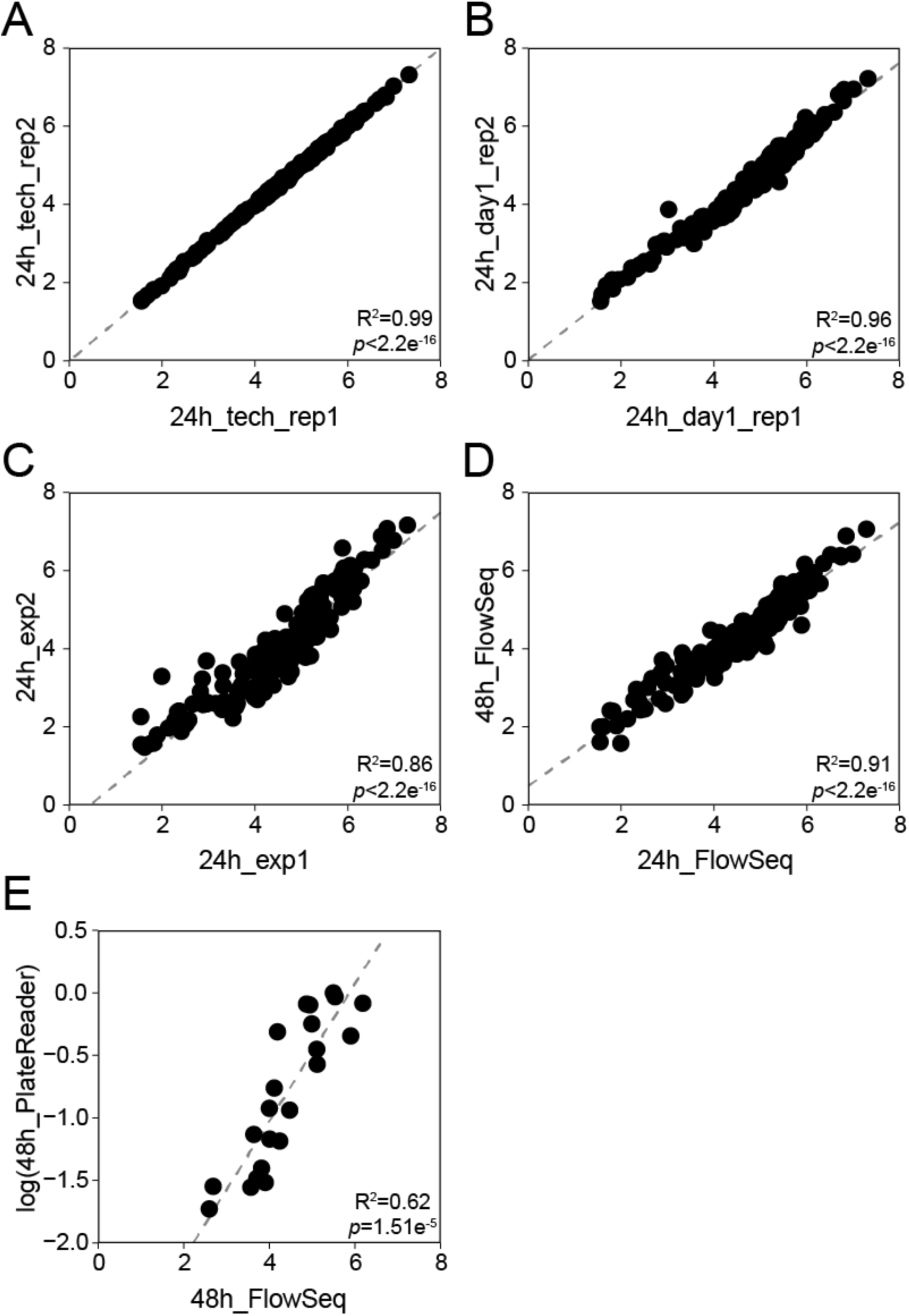
Reproducibility of Flow-seq experiments in HeLa cells (unspliced GFP variants). (A) Re-sequencing of the same amplicon-library. (B-C) Replicate Flow-seq experiments performed on the same day (B) or different days (C). (D) Flow-Seq experiments performed on the same pool of cells, 24h and 48h after the induction of GFP expression. (E) Correlation between fluorescence measurements of 22 GFP variants obtained spectrofluorometry of transiently transfected HeLa cells and by Flow-Seq of HeLa GFP pool cell line.

**Supplementary Figure 5.**
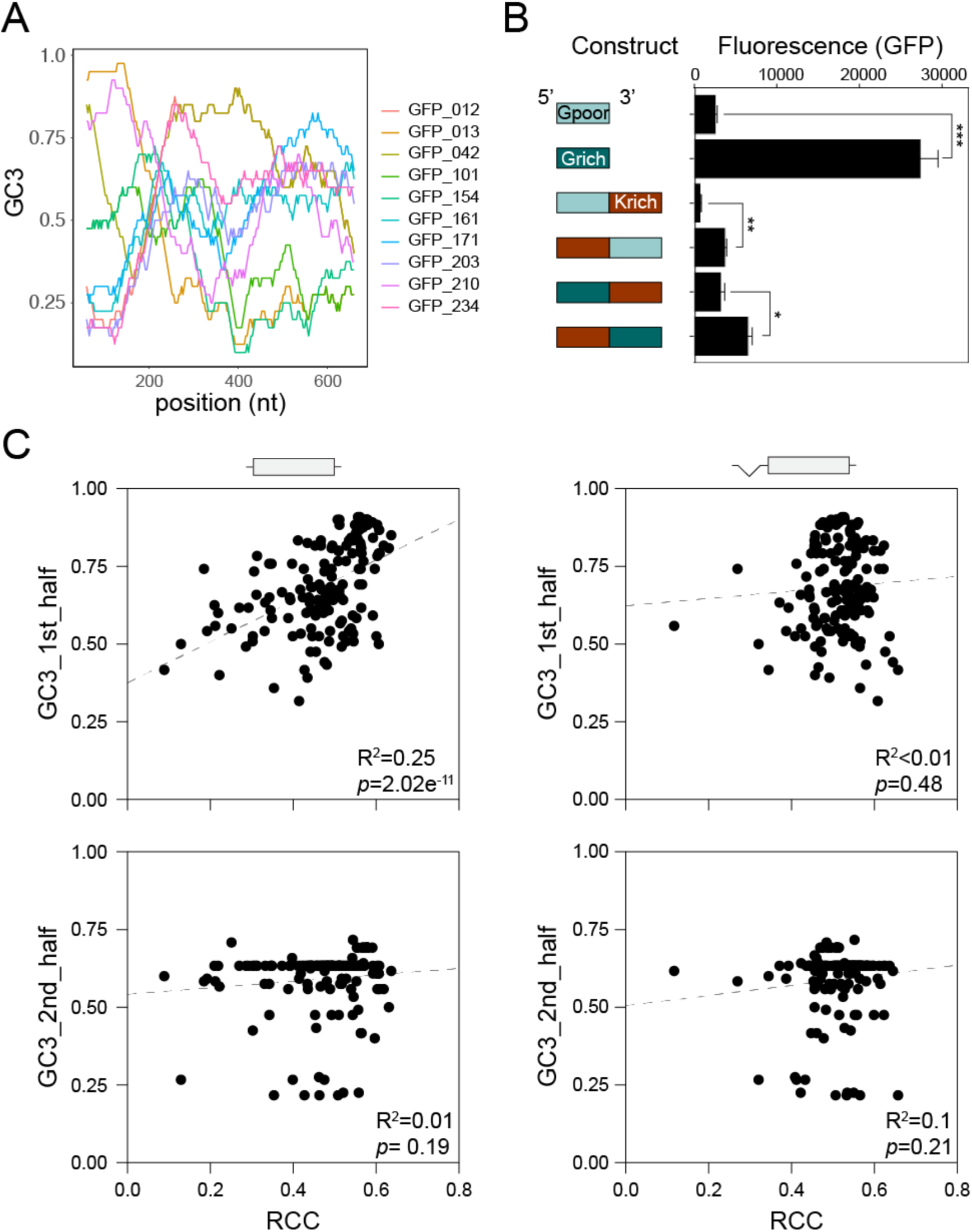
Position-specific effects of GC content on expression. (A) Sliding window analysis of GC3 content in selected GFP variants used in the pooled amplicon sequencing experiments. (B) Protein measurements of translational fusion constructs between GC-poor (33% GC3, Gpoor) and GC-rich (97% GC3, Grich) variants of GFP with a GC-rich variant of mKate2 (85% GC3, Krich), upon transient transfection into HeLa cells. Data represent the mean of 3 replicates + SEM. (C) Correlations between the GC3 content in the 1st (nt 1-360) and 2nd (nt 361-720) halves of GFP variants and their relative cytoplasmic mRNA concentrations.

**Supplementary Figure 6.**
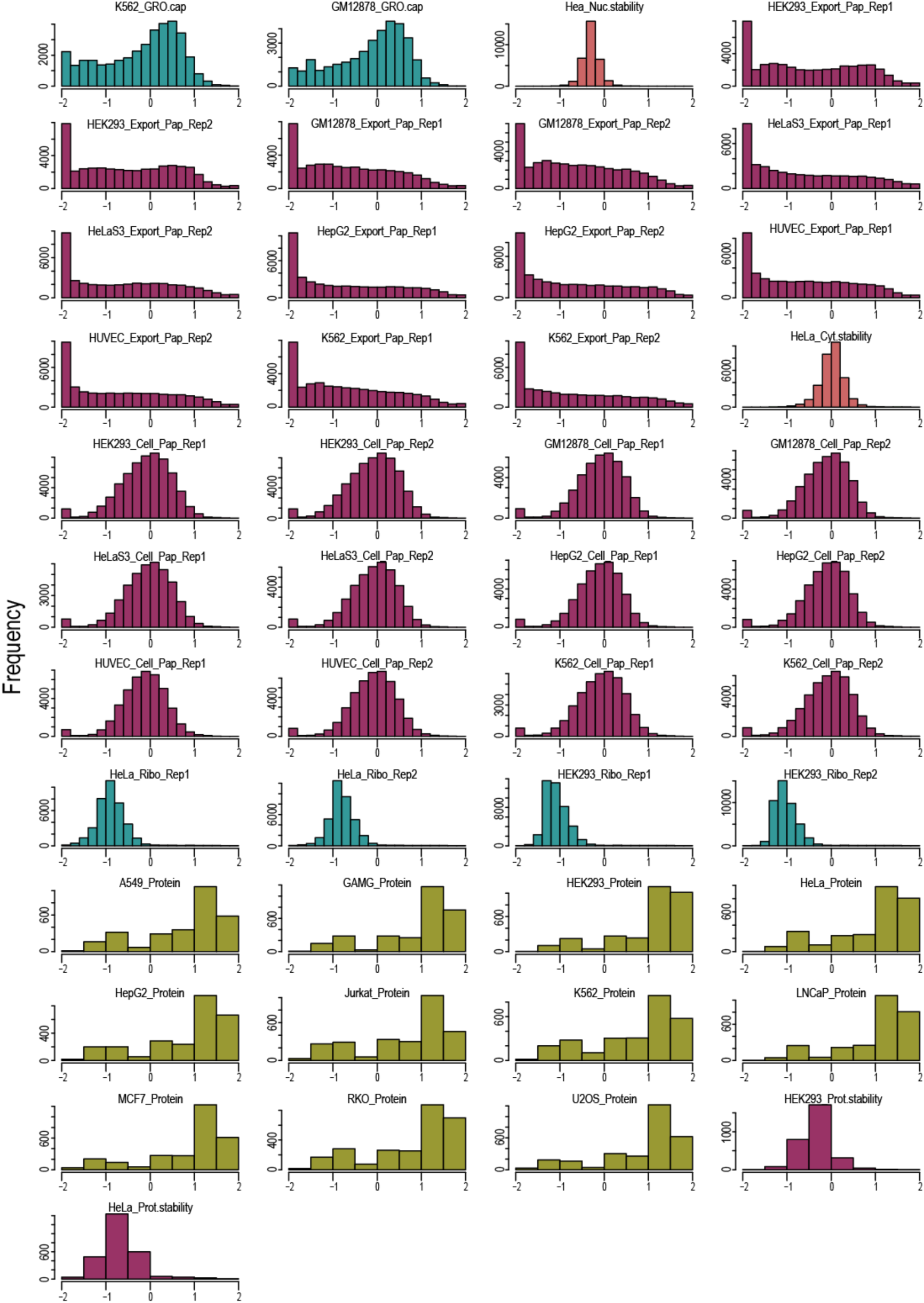
Distribution of RNA and protein expression data used in regression modelling. Human RNA and protein expression data were extracted from various databases, filtered and normalized as described in Supplementary Table 1 and in the Methods section. The histograms show the distributions of preprocessed expression measurements.

**Supplementary Table 1.**
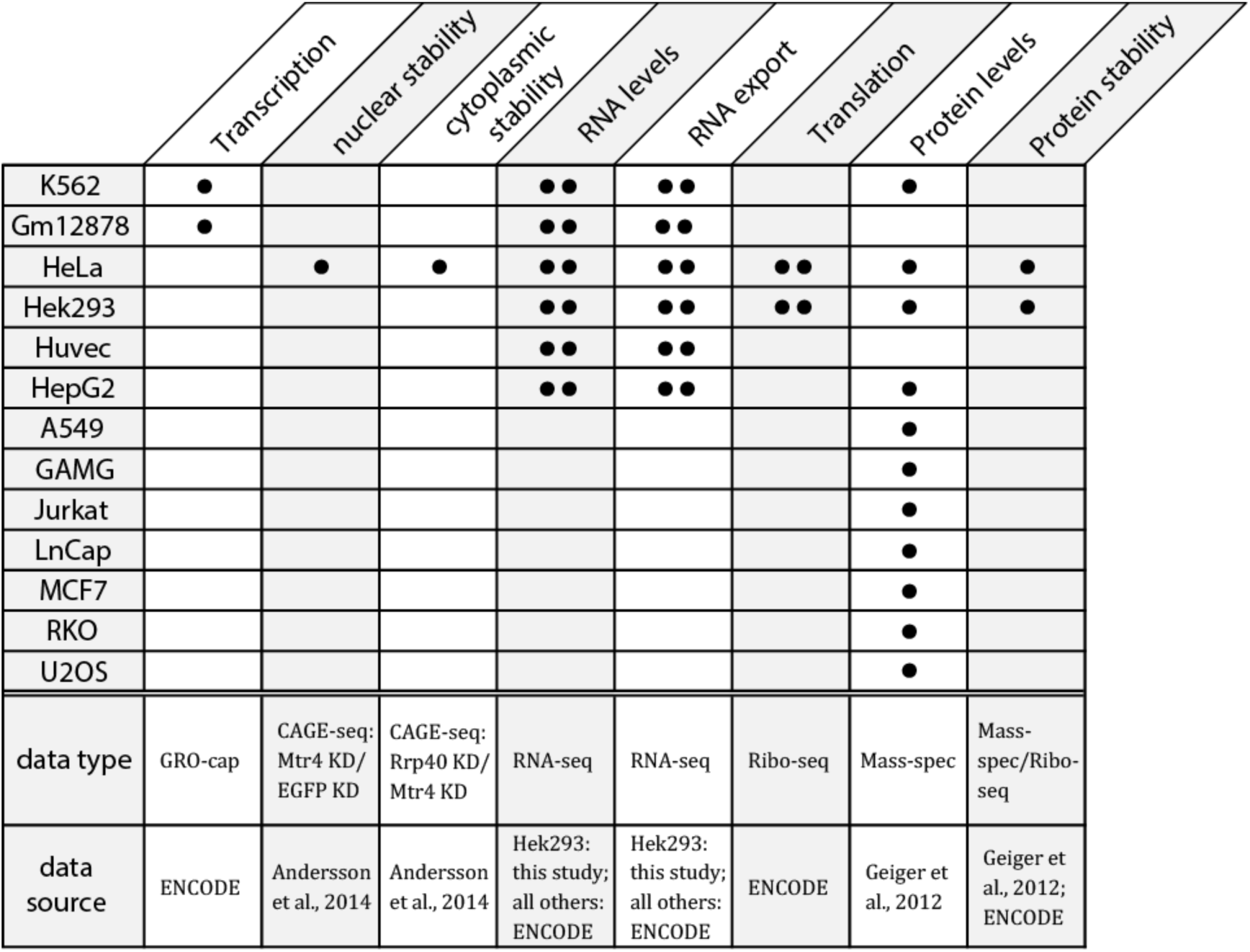
Sources of human gene expression data. The cellular process to be quantified is indicated above the table, and the experimental techniques and data sources are indicated below. Each dot indicates an experimental replicate measurement.

|  | Transcription | nuclear stability | cytoplasmic stability | RNA levels | RNA export | Translation | Protein levels | Protein stability |
| --- | --- | --- | --- | --- | --- | --- | --- | --- |
| K562 | • |  |  | •• | •• |  | • |  |
| Gm12878 | • |  |  | •• | •• |  |  |  |
| HeLa |  | • | • | •• | •• | •• | • | • |
| Hek293 |  |  |  | •• | •• | •• | • | • |
| Huvec |  |  |  | •• | •• |  |  |  |
| HepG2 |  |  |  | •• | •• |  | • |  |
| A549 |  |  |  |  |  |  | • |  |
| GAMG |  |  |  |  |  |  | • |  |
| Jurkat |  |  |  |  |  |  | • |  |
| LnCap |  |  |  |  |  |  | • |  |
| MCF7 |  |  |  |  |  |  | • |  |
| RKO |  |  |  |  |  |  | • |  |
| U2OS |  |  |  |  |  |  | • |  |
| data type | GRO-cap | CAGE-seq: Mtr4 KD/ EGFP KD | CAGE-seq: Rrp40 KD/ Mtr4 KD | RNA-seq | RNA-seq | Ribo-seq | Mass-spec | Mass-spec/Ribo-seq |
| data source | ENCODE | Andersson et al., 2014 | Andersson et al., 2014 | Hek293: this study; all others: ENCODE | Hek293: this study; all others: ENCODE | ENCODE | Geiger et al., 2012 | Geiger et al., 2012; ENCODE |

**Supplementary Table 2.**
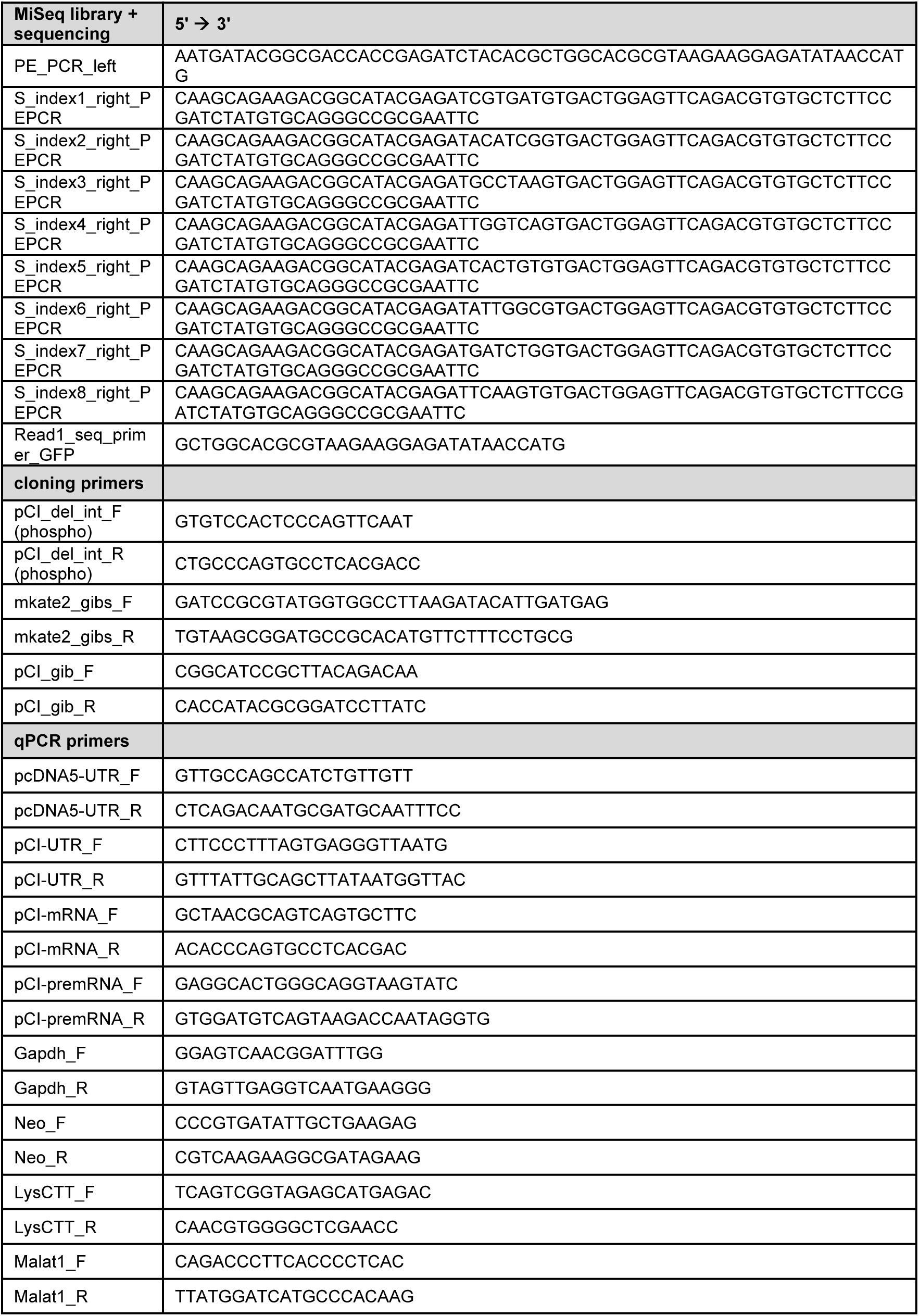
List of primers used.

